# Single-cell decoding of human islet cell type-specific alterations in type 2 diabetes reveals converging genetic- and state-driven β-cell gene expression defects

**DOI:** 10.1101/2025.01.17.633590

**Authors:** Khushdeep Bandesh, Efthymios Motakis, Siddhi Nargund, Romy Kursawe, Vijay Selvam, Redwan M Bhuiyan, Giray Naim Eryilmaz, Sai Nivedita Krishnan, Cassandra N. Spracklen, Duygu Ucar, Michael L. Stitzel

## Abstract

Pancreatic islets maintain glucose homeostasis through coordinated action of their constituent endocrine and affiliate cell types and are central to type 2 diabetes (T2D) genetics and pathophysiology. Our understanding of robust human islet cell type-specific alterations in T2D remains limited. Here, we report comprehensive single cell transcriptome profiling of 245,878 human islet cells from a 48-donor cohort spanning non-diabetic (ND), pre-diabetic (PD), and T2D states, identifying 14 distinct cell types detected in every donor from each glycemic state. Cohort analysis reveals ∼25-30% loss of functional beta cell mass in T2D vs. ND or PD donors resulting from (1) reduced total beta cell numbers/proportions and (2) reciprocal loss of ‘high function’ and gain of senescent β-cell subpopulations. We identify in T2D β-cells 511 differentially expressed genes (DEGs), including new (66.5%) and validated genes (e.g., *FXYD2, SLC2A2, SYT1*), and significant neuronal transmission and vitamin A metabolism pathway alterations. Importantly, we demonstrate newly identified DEG roles in human β-cell viability and/or insulin secretion and link 47 DEGs to diabetes-relevant phenotypes in knockout mice, implicating them as potential causal islet dysfunction genes. Additionally, we nominate as candidate T2D causal genes and therapeutic targets 27 DEGs for which T2D genetic risk variants (GWAS SNPs) and pathophysiology (T2D vs. ND) exert concordant expression effects. We provide this freely accessible atlas for data exploration, analysis, and hypothesis testing. Together, this study provides new genomic resources for and insights into T2D pathophysiology and human islet dysfunction.

## Introduction

Pancreatic islets are primary mediators of type 2 diabetes (T2D) genetic risk and pathophysiology, driving insulin secretion defects^1,2,3^. They comprise multiple cell types, including insulin-secreting beta (β) cells, glucagon-secreting alpha (α) cells, somatostatin-secreting delta (δ) cells, pancreatic polypeptide-secreting gamma (γ) cells and ghrelin-secreting epsilon (ε) cells, that collectively determine islet functional output^4^. In humans, they are intermingled throughout the islet, ensuring equal access to vasculature to sense and respond to fluctuating glucose levels^5^. Growing evidence shows that α- and δ-cell signals regulate β-cell function to ensure proper insulin secretion dynamics^5^. Robust assessment of islet cell type-specific gene expression programs and their regulation in pathologic states is crucial to define and understand pancreatic dysfunction in T2D. However, donor variability, modest sample size, and/or a relatively small number of cells sampled per individual have limited power to detect robust, reproducible differences^6–10^. To address this challenge, we completed single cell transcriptome profiling and analysis of pancreatic islets obtained from a cohort of 48 non-diabetic (ND), pre-diabetic (PD), and type 2 diabetic (T2D) donors. We identified robust T2D-associated differences in islet cell type composition and gene expression and nominated 92 β-cell differentially expressed genes (DEGs) for putative causal roles in islet dysfunction using complementary experimental, physiologic, and genetic approaches and analyses.

## Results

### Comprehensive human islet single-cell transcriptome atlas spanning non-diabetic, pre-diabetic, and type 2 diabetic states

To build a comprehensive, representative atlas of human islet transcriptomes, we completed single cell RNA sequencing (scRNA-seq) in human pancreatic islets obtained through the Integrated Islet Distribution Program (IIDP) from a diverse cohort of 48 American cadaveric organ donors, representing European, Hispanic, and African American self-reported ancestries and independent from those in the Human Pancreas Analysis Program (HPAP)^6,11^ (**Supplementary Table 1**). The cohort included 17 diagnosed T2D (mean HbA1c = 7.6%), 14 PD (mean HbA1c = 5.9%; designated based on American Diabetes Association (ADA) prediabetes criteria (5.7% ≤ HbA1c ≤ 6.4%)^12^), and 17 ND (mean HbA1c = 5.2%) donors (**Figure 1a**). ND control donor samples were selected to be as similar as possible to the T2D cases with respect to age, sex, BMI, and self-reported ancestry. HbA1c levels differed significantly between groups, but age and BMI were similar (**Figure 1b**, Games-Howell post-hoc test) and sex and ancestry distributions were comparable (**Supplementary Table 1**). Donor islets were dissociated into single-cell suspensions, which were captured and sequenced to a median sequencing depth of 13,400 UMIs using droplet-based scRNA-seq (10X Genomics). In total, 245,878 cells passed stringent quality control (see Methods), with an average of 5,122 high quality single cells per donor (**Supplementary Figures 1a** and **1b** and **Supplementary Table 2**). After batch correction (**Supplementary Figure 1c**), unsupervised clustering based on expression of the 2,000 most variable genes among these single cell transcriptomes identified 14 distinct cell types corresponding to endocrine (α, proliferating α, β, δ, γ, and ε), exocrine (acinar and ductal), stellate/activated stellate, endothelial, glial (Schwann), and resident (mast) and infiltrating immune cell types across ND, PD, and T2D donors (**Figure 1c** and **Supplementary Figure 2**).

**Figure 1:**
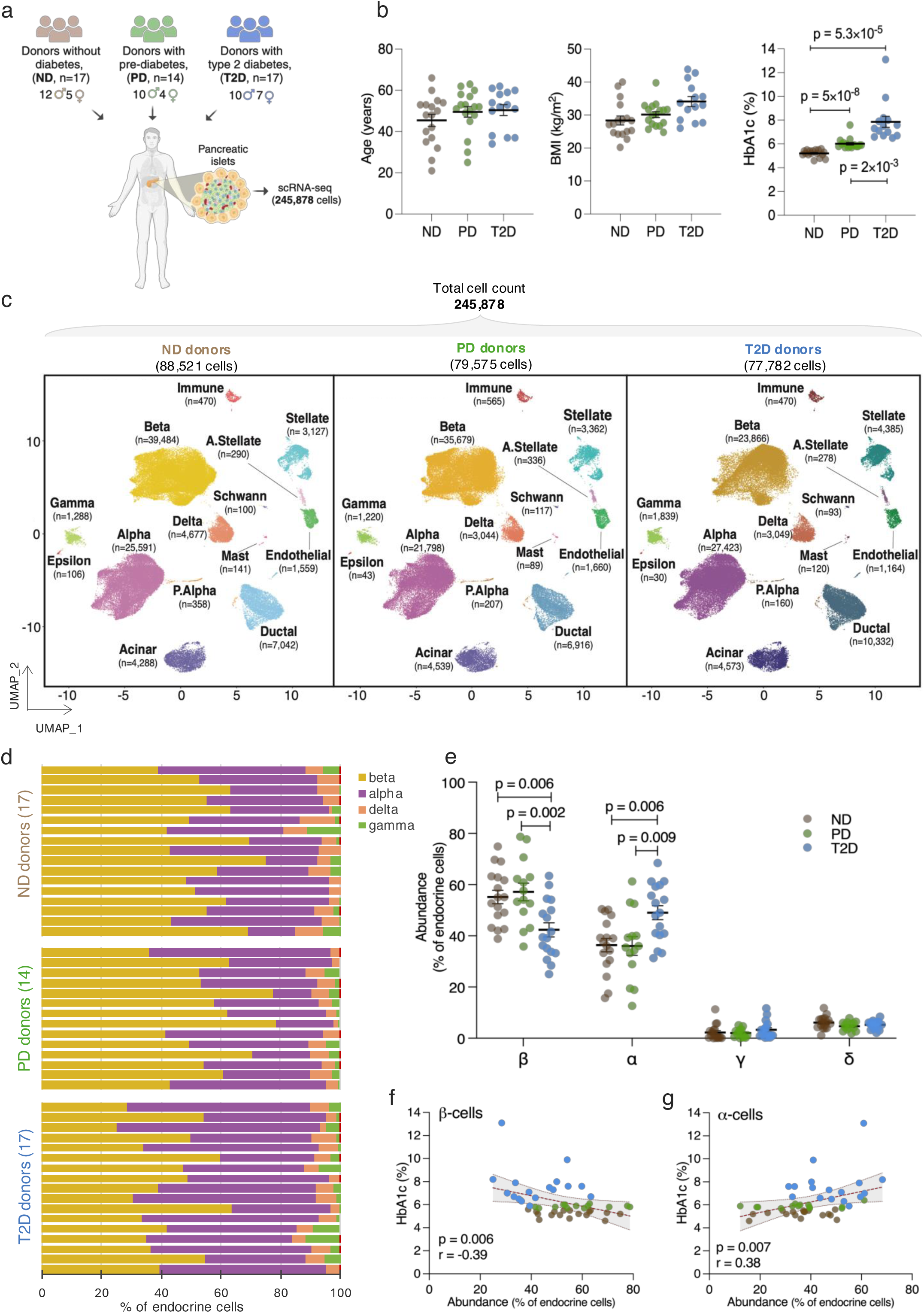
Human pancreatic islet single-cell transcriptomes from 48-donor cohort reveal cell type proportion variability in T2D donors. **(a)** Human pancreatic islets from 48 cadaveric donors -- 17 with diagnosed type 2 diabetes (T2D), 14 with HbA1c-based prediabetes (PD; HbA1c 5.7%-6.4%), and 17 without diabetes (ND) -- were dissociated into single cells and profiled using droplet-based scRNA-seq to obtain 245,878 high quality islet single cell transcriptomes. **(b)** Comparison of age, body mass index (BMI, a measure of obesity), and glycated hemoglobin (HbA1c) between T2D, PD, and ND individuals in the cohort. Each dot represents an individual donor. The black line and error bars represent the mean ± standard error of the mean. Significant differences (p<0.05, Games-Howell post-hoc test) are reported. **(c)** Uniform Manifold Approximation and Projection (UMAP) plots displaying unsupervised clustering of 245,878 cells in ND (left), PD (middle), and T2D (right) donors reveals 14 distinct cell types based on the expression of the 2000 most variable genes across the cells. n=number of single cell transcriptomes obtained for each cell type. **(d)** Relative percentages of various endocrine cell types, shown in per-donor stacked bar plots across the glycemic states, indicate remarkably fewer β-cells in islets from T2D (bottom) vs. PD (middle) or ND (top) donors. **(e)** Relative abundance of α-, β-, δ-, and γ-cells in ND, PD, or T2D donors. Dots represent percentage of endocrine cells detected for each donor. Epsilon (ε) cells were rare (0.09%) in all donors and omitted from comparison. The black line and error bars represent the mean ± standard error of the mean. P-values were calculated using Tukey’s honest significance test. Significant differences (p<0.05) are indicated. **(f, g)** Spearman correlations between HbA1c levels (y-axis) and relative β-cell **(f**, x-axis) or α-cell **(g**, x-axis) abundance for all cohort donors (n=48). Bands enclosing the linear regression line represent 99% confidence intervals. Dots represent individual donors colored as in panel (**a**) based on their glycemic status.

We defined robust signature genes—those specific to each islet cell type and detected across donors irrespective of their glycemic status—by aggregating and comparing per-donor single cell transcriptomes of individual cell types. In addition to classic hormone-encoding marker genes such as *INS*, *GCG*, *SST*, *PPY*, and *GHRL*, we identified 270 α-, 272 β-, 173 δ-, 130 γ-, and 194 ε-cell signature genes exhibiting ≥ 8-fold expression differences at a false discovery rate (FDR) <5% in one-vs-all ANOVA comparisons (**Supplementary Figure 2; Supplementary Table 3**). Functional processes enriched among these signature genes included G protein-coupled receptor signaling and amino acid transport (α-cells); insulin secretion, regulation of membrane potential, and neuronal transmission (β-cells); gamma-aminobutyric acid signaling/synaptic transmission and synapse assembly (δ-cells); G protein-coupled receptor signaling, neuropeptide signaling and regulation of cation channel activity (γ-cells); and regulation of lipoprotein lipase activity (ε-cells) (**Supplementary Table 4**). Examining sex differences, we compared gene expression between males and females across all states combined, identifying 112 α-, 64 β-, and 45 δ-cell DEGs by sex, 27 of which were shared across these three cell types (**Supplementary Table 5**). 26/27 sex-specific DEGs were on X or Y chromosomes (**Supplementary Table 5**) and were not significantly enriched for any common processes or pathways.

### Significant β-cell loss in T2D vs. PD, ND donors

After establishing the islet cell types and their robust expression signatures, we assessed T2D-associated alterations in islet cell type composition by comparing cell type counts obtained from each islet donor between ND, PD, and T2D states (**Supplementary Table 6**). The distribution of each endocrine cell type relative to the total number of endocrine cells confirmed substantial inter-donor heterogeneity within each state (**Figure 1d**)^7,8,13^. In T2D islets, β-cell/endocrine proportions were 13-15% lower (mean β-cell %=42.2±11.3) than those in ND (mean β-cell %=55.2±10.7, p=0.006) or PD (mean β-cell %=57.2±12.9, p=0.002, **Figure 1e**, ANOVA followed by Tukey’s honest significance test) donor islets. α-cell proportions were correspondingly higher in T2D islets (48.7±11.2% vs. 35.8±10.5% in ND (p=0.006) or vs. 35.7±13.6% in PD (p=0.009)). Relative proportions of δ- and γ-cells were similar between groups.

A portion of the islets from 30/48 donors in this single-cell transcriptome cohort were also characterized by the IIDP Human Islet Phenotyping Program (HIPP) (https://iidp.coh.org/Resources-Offered/HIPP), which reported immunofluorescence-based estimates of their islet cell type composition (**Supplementary Table 6**). Cell proportions calculated from per-donor scRNA-seq profiles and HIPP for these samples were correlated (**Supplementary Figure 3**; r=0.62 for ND, 0.70 for PD, and 0.73 for T2D for β-cells, and r=0.65 for ND, 0.81 for PD, and 0.74 for T2D for α-cells), suggesting that the scRNA-seq-determined cell proportion differences were not due to cell loss during sample processing or single cell capture. Collective analysis of the full cohort revealed a significant inverse correlation between reduced β-cell proportions and elevated HbA1c levels (**Figure 1f**, Spearman’s r=-0.39; p=0.006), consistent with reported inverse associations between HOMA-β (an index reflecting β-cell function) and HbA1c levels^14^, while α-cell expansion also correlated with elevated HbA1c (**Figure 1g**, Spearman’s r=0.38; p=0.007).

### Cell type-specific gene expression differences in T2D islets

Next, we sought to identify robust cell type-specific gene expression differences between ND, PD, and T2D individuals by aggregating each individual’s scRNA-seq profiles per cell type into “pseudobulk” gene expression profiles and comparing them (Methods). Surprisingly, we identified only 3 α- (*GGT6* and *GALNT13* (T2D vs. ND); *AC104770.1* (PD vs. ND)), 2 δ- (*TMEM190* (T2D vs. ND); *CTRB1* (PD vs. ND)), and 6 γ- (*MEG3*, *MEG8*, *LPO* and *CA2* (T2D vs. ND); *GAL* and *AKR1C3* (T2D vs. PD) cell-specific DEGs between the 3 glycemic states at FDR< 5% (**Supplementary Tables 7, 8 and 9**). In striking contrast, we discovered 746 β-cell DEGs in T2D vs. ND donors at FDR<5% (**Figure 2a** and **Supplementary Table 7**), approximately 10 times the number reported in recent HPAP cohort-based analyses^6^. 511 β-cell DEGs exhibited a fold change (FC) ≥ 50% (i.e. log_2_FC ±0.585), 316 or 195 of which were up- or down-regulated, respectively. We replicated 171 T2D DEGs reported in previous whole islet or cell type-specific studies^6–9,15–21^, including *FXYD2*, *SLC2A2*, *SCN9A*, *PAX5*, *DGKB*, *IRS1* and *SYT1*. Importantly, two-thirds of detected β-cell DEGs were previously unreported (n=340; **Supplementary Figure 4a-d** and **Supplementary Table 10**). Because sample sizes were insufficient for meaningful analyses or interpretation, we did not assess sex-specific differences in this cohort.

**Figure 2:**
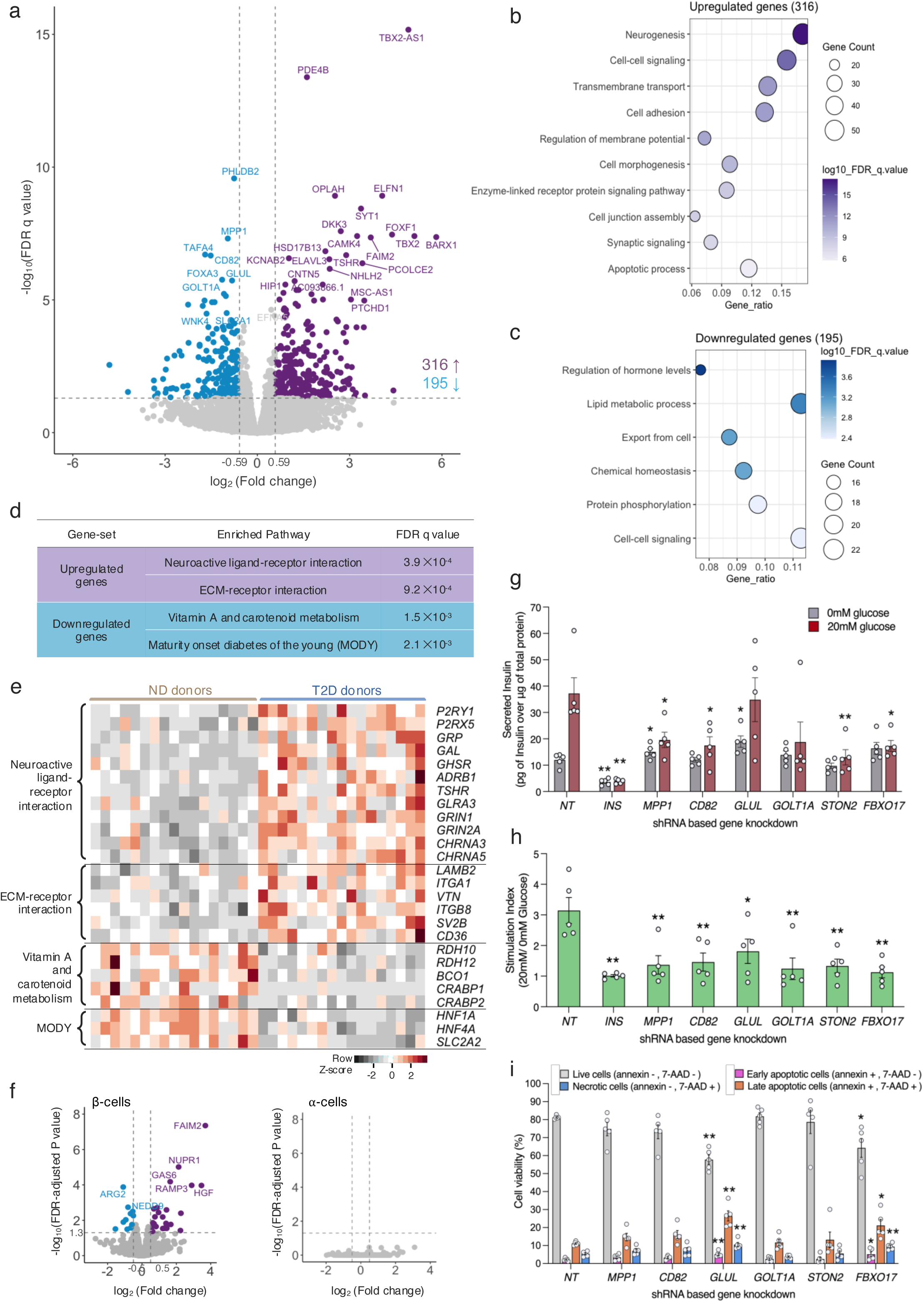
Differentially expressed genes in T2D vs. ND β-cells. **(a)** Volcano plot of differentially expressed genes in T2D vs. ND β-cells. Each dot denotes a gene. 511 genes with significant differences in expression at false discovery rate (FDR) < 5% and fold change ≥ 50% are colored blue (T2D-downregulated) or purple (T2D-upregulated); gray dots denote those with comparable expression in T2D and ND β-cells. (**b,c**) Gene set enrichment analysis (GSEA) for differentially expressed genes using the molecular signatures database (MSigDB, BROAD Institute). Enriched non-redundant processes with FDR q<0.05 are shown for upregulated **(b)** and downregulated **(c)** gene sets. Number of genes in each functional term is provided as a gene ratio relative to total number of tested upregulated (n=316) or downregulated (n=195) T2D β-cell genes. **(d)** KEGG- and Wiki-pathways enriched in up- vs. downregulated genes. **(e)** Heatmap of scaled expression of genes comprising primary enriched molecular pathways identified **(d)** in T2D vs. ND donors. **(f)** Volcano plots showing differentially expressed cell death genes in β- (left) or α- (right) cells from T2D vs. ND donor islets. **(g)** Basal (gray; 0mM glucose) or glucose-stimulated (red; 20 mM glucose) insulin secretion in human EndoC-βH3 cells following shRNA-mediated knockdown of selected target genes or a non-targeting control sequence (NT). Data represent mean ± standard error of the means (s.e.m.) from 5 independent experiments, each represented by dots. Significance was calculated relative to corresponding conditions for NT control cells using unpaired Student’s t-test where *p < 0.05 and **p < 0.01. **(h)** Stimulation Index (SI) for shRNA gene knockdowns, calculated from high vs. low glucose insulin secretion measured in panel (**g**). **(i)** EndoC-βH3 cell viability (assessed via Annexin V and 7-AAD staining) showing relative percentages of viable, early- or late-apoptotic, or necrotic cells after shRNA knockdown of selected genes or NT control. Data represent mean ± s.e.m. from 5 independent flow cytometry experiments, each represented by dots. Significance was assessed relative to corresponding measures in NT cells; *p < 0.05 and **p < 0.01; Student’s t-test.

In the full cohort, the five T2D β-cell protein-coding genes with the most significant FDR were those with neuronal functions^22^, mediating central nervous system effects of therapeutic agents (*PDE4B*) or acetyl-choline receptor aggregation in postsynaptic membranes (*PHLDB2*), catalyzing proline conversion to the major excitatory neurotransmitter glutamate (*OPLAH*), regulating postsynaptic neural circuit dynamics (*ELFN1*), or sensing calcium for neurotransmitter release (*SYT1*). 4/5 DEGs (excluding *PHLDB2*) were detected in both male and female T2D donors when analyzed separately. More broadly, 17.1% of T2D-upregulated β-cell genes were enriched for the biological term ‘neurogenesis’ (FDR p value, q = 6.0 ×10^−18^), followed by ‘cell-cell signaling’ (q = 1.0 ×10^−14^), ‘cell adhesion’ (q = 5.1 ×10^−12^), and ‘regulation of membrane potential’ (q = 1.7 ×10^−11^) (**Figure 2b** and **Supplementary Table 11**). More than 15% of the newly identified T2D-upregulated β-cell genes were associated with the cellular component ‘synapse’ (q = 1.1 ×10^−7^). Conversely, downregulated T2D β-cell genes were enriched for the processes ‘regulation of hormone levels’ (q = 1.2 ×10^−4^), ‘lipid metabolism’ (q = 4.3 ×10^−4^), and ‘cell-cell signaling’ (q = 4.2 ×10^−3^, **Figure 2c**). Pathway enrichment analyses (MSigDB, https://www.gsea-msigdb.org/gsea/msigdb) revealed ‘neuroactive ligand-receptor interaction’ (q = 3.9 ×10^−4^) and ‘vitamin A and carotenoid metabolism’ (q = 1.5 ×10^−3^) as the primary molecular pathways associated with the up- and downregulated DEGs, respectively (**Figure 2d** and **Supplementary Table 11**).

The enriched neuroactive ligand-receptor interaction pathway included genes encoding glutamate (*GRIN1*, *GRIN2A*), acetylcholine (*CHRNA3*, *CHRNA5*), and norepinephrine (*ADRB1*) neurotransmitter receptors; neuroendocrine peptides (*GRP*, *GAL*); ATP receptors (*P2RY1*, *P2RX5*); receptors for hormones with known insulin suppressive effects (*GHSR*, *TSHR*); and a ligand-gated ion channel (*GLRA3*) (**Figure 2e** and **Supplementary Figure 5a**). Pancreatic islets are densely innervated by autonomic nerves^23^, and the preganglionic sympathetic and parasympathetic neurons innervating islets primarily release acetylcholine^24^. Additionally, postganglionic sympathetic neurons release norepinephrine, glutamate, and galanin (encoded by *GAL*) while postganglionic parasympathetic neuron fibers release gastric releasing peptide (encoded by *GRP*)^26^. Earlier studies in nerve stimulation have demonstrated the ability of neural signals to override the effects of circulating glucose^25,26^. Individuals with T2D exhibit increased islet innervation, possibly as a compensatory mechanism by which the nervous system tries to preserve or augment islet function under metabolic stress^23^. Presence of multiple neuroreceptors on β-cells indicate that they can directly sense and respond to neural signals. Altered expression of these neuroreceptors or neuropeptides within β-cells from individuals with T2D suggests disrupted neuroendocrine regulation of insulin secretion as a pathophysiologic aspect of pancreatic dysfunction that is not yet fully understood in T2D. In addition, our data highlight underappreciated roles for aberrant purinergic signaling, involving upregulated ATP receptor genes (*P2RY1*, *P2RX5*), in islet dysfunction and T2D. Notably, insulin secretory vesicles contain ATP and ADP molecules, which are co-released with insulin during glucose-stimulated exocytosis^27^. These secreted purine adenosines act as extracellular signaling mediators that activate two types of purinergic P2 receptors on the β-cell membrane—P2X (ligand-gated cation channels) and P2Y (G-protein coupled channels)—whose autocrine effects amplify glucose-induced calcium [Ca^2+^] responses in β-cells^28^.

Downregulation of multiple genes encoding key proteins in vitamin A metabolism— retinol dehydrogenases (*RDH10*, *RDH12*), β-carotene oxygenase (*BCO1*, which generates the vitamin A precursor, β-carotene), and cellular retinoic acid binding proteins (*CRABP1* and *CRABP2*) (**Supplementary Figure 5b**)—was another hallmark of T2D β-cells. Vitamin A metabolites (retinoic acid, RA) regulate gene expression by activating transcription factors (e.g., HNF4A, retinoid A receptor nuclear receptor (RAR)) that bind to RA-response elements in target genes. Dietary deficiency of vitamin A has been linked with hyperglycemia, mirroring reduced vitamin A levels in the pancreas^29^. Retinoids are regulators of apoptosis^30,31^, and vitamin A deprivation has been shown to decrease β-cell mass^29^. To test the apoptotic impact of compromised vitamin A metabolism in T2D, we investigated the expression of 1,159 genes encoding proteins associated with cell death in the human protein atlas (HPA, https://www.proteinatlas.org)^32^. T2D vs ND β-cells exhibited differential expression of several of these genes including induction of *FAIM2*, *NUPR1*, *GAS6*, *HGF*, and *RAMP3* and reduction of *ARG2* and *NEDD9*, none of which were modulated in T2D α-cells (**Figure 2f**). Many of these genes harbor RA response elements (**Supplementary Figure 6**) and are anti-apoptotic, which suggests a compensatory response to vitamin A deficiency and related increase in β-cell apoptosis.

T2D β-cell DEGs included multiple genes not previously linked to islet β-cell dysfunction or T2D. We selected a subset of these new causal candidates for functional validation based on their suspected roles in T2D and informed by their general cellular functions. Specifically, we targeted *MPP1*, *CD82*, *GLUL,* and *GOLT1A* (4/10 most downregulated genes), *STON2* (an adaptor protein involved in recycling synaptic vesicles for neurotransmission^22^), and *FBXO17* (a regulator of Akt signaling pathway^22^). To evaluate their role in insulin secretion, we completed shRNA knockdown and assessed effects on glucose-stimulated insulin secretion (GSIS) assays in human EndoC-βH3 β-cells. Compared to non-targeting (NT) shRNA control cells, knockdown of all targets altered basal insulin secretion, GSIS, or both. *MPP1* and *GLUL* knockdown increased basal insulin secretion (**Figure 2g** and **Supplementary Figure 7**), a feature associated with islet dysfunction in T2D^33^. When stimulated with 20mM glucose, *MPP1*, *CD82*, *STON2,* and *FBXO17* knockdown cells exhibited blunted insulin secretory responses. Assessment of the stimulation index revealed impaired GSIS for all 6 genes following knockdown (**Figure 2h**). Furthermore, Annexin V/7-AAD staining indicated that *GLUL* and *FBXO17* knockdown markedly and modestly decreased cell viability, respectively, with consequent increases in early apoptotic, late apoptotic, and necrotic cells relative to shControl cells (**Figure 2i**). *GLUL* encodes glutamine synthetase, which catalyzes conversion of glutamate and ammonia to glutamine. Matschinksy and colleagues previously demonstrated roles for glutamine in both amino acid- and GSIS^34^, but *GLUL* has not been directly linked to T2D. Elevated plasma glutamate levels and lower plasma glutamine levels associate with increased T2D incidence^35^, and GLUL uniquely executes their biochemical conversion^22^. Considering its pivotal role in these metabolic pathways and the evidence from previous studies, investigating *GLUL* as a potential target in T2D research holds considerable promise.

### Convergent T2D genetic and pathophysiologic effects on expression nominate T2D causal genes

In addition to directly testing selected DEG effects on human beta-cell viability and function, we sought to nominate additional DEGs contributing to, rather than a consequence of, T2D physiology based on links to glucose homeostasis and T2D genetics. Interestingly, genes harboring inactivating monogenic diabetes mutations, such as *HNF1A*, *HNF4A,* and *SLC2A2*^36^, were significantly downregulated in T2D donor β-cells (**Figure 2e**). Given these concordant down-regulation/loss-of-function effects, we sought to identify T2D β-cell DEGs for which T2D-associated risk alleles exerted concordant gene expression effects. We compiled a list of 39,972 T2D-associated index and linked proxy genetic variants (all-ancestry or ancestry-specific LD r^2^≥0.80 in 1000Genomes Phase 3 data) reported in genome-wide association study (GWAS) meta-analyses from T2DGGI^37^, DIAMANTE^38^, MVP^39^, and AGEN^40^ (**Supplementary Table 12**). We queried islet expression QTL association results (p < 0.05) from the TIGER consortium^41^ and identified 461 T2D variants (representing 41 loci) associated with altered islet expression of 25 upregulated and 16 downregulated genes in T2D β-cells. T2D genetic and environmental effects were concordant (i.e., T2D risk allele altered islet gene expression in the same direction as T2D vs. ND β-cell differential expression) for 27 genes (**Figure 3a**, red and blue; **Supplementary Table 13**), including *DGKB, ST6GAL1,* and *STARD10* reported as colocalized T2D genetic and islet eQTL association signals showing directionally consistent impact on gene expression^41,42,43^.

**Figure 3:**
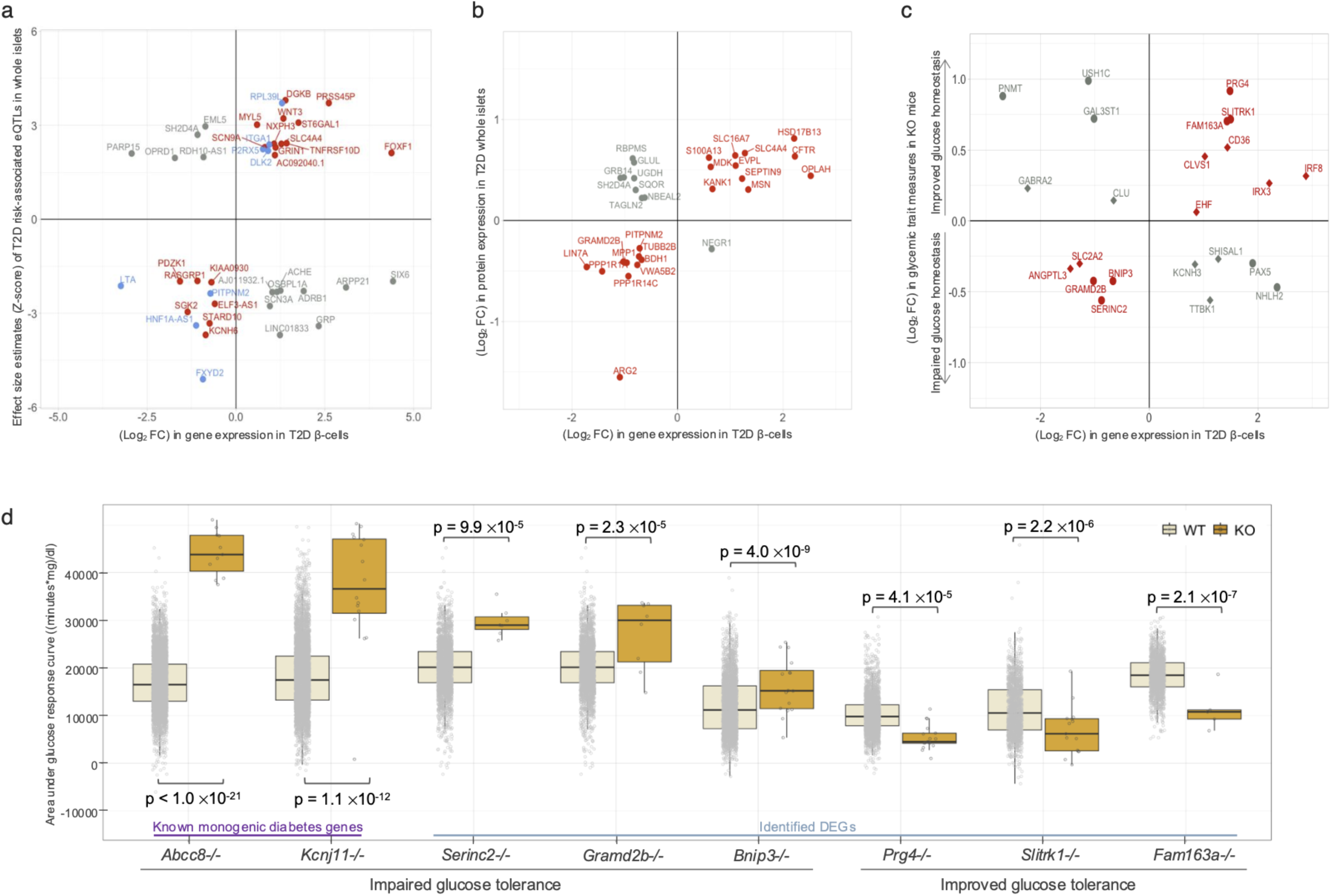
Integrated multimodal analyses prioritize T2D β-cell differentially expressed genes as candidate causal/driver genes. **(a)** Comparison of islet eQTL effect sizes from TIGER consortium^41^ (y-axis) vs. fold-change in gene expression for T2D β-cell differentially expressed genes (DEGs, x-axis) from this study. Red denotes genes with consistent T2D genetic and disease state effects on expression. Upper right quadrant genes are concordantly upregulated; lower left quadrant genes are concordantly downregulated. Gray denotes genes with opposite islet eQTL vs. T2D β-cell differential expression effects. Blue denotes genes with multiple T2D genetic association signals exhibiting concordant and discordant islet eQTL effects. **(b)** Comparison of T2D differentially abundant proteins in Humanislets^56^ database (y-axis) and T2D β-cell DEGs (x-axis). Red and gray denote genes with concordant or discordant β-cell RNA and islet protein level differences in T2D vs. ND individuals, respectively. **(c)** T2D β-cell DEGs significantly associated with various glycemic phenotypes in whole-body knockout (KO) mice from the IMPC^62^ consortium data. Log_2_ fold change (FC) gene expression in T2D vs. ND β-cells (x-axis) compared to Log_2_ fold change in trait measure in KO mice vs. wild-type mice (y-axis). Circles or diamonds distinguish glucose tolerance (area under glucose response curve and initial response to glucose challenge) vs. fasting glucose phenotypes, respectively. Red denotes genes with KO glycemic phenotypes consistent with T2D physiology while gray genes indicate a T2D-misaligned defect. **(d)** IMPC glucose homeostasis phenotypes of known diabetes gene (*Abcc8* and *Kcnj11*) and T2D β-cell DEG KO mice compared to wildtype controls. Glucose tolerance measured by initial response to glucose challenge and/or area under the glucose response curve from an intraperitoneal glucose tolerance test (IPGTT). See **Supplementary Table 14** for individual measures.

For nineteen T2D β-cell DEGs, we identified a single T2D GWAS variant for which the T2D risk allele altered expression in the same direction. For example, the T2D risk allele ‘C’ of intergenic variant rs35825770 (OR = 1.02, p = 3.5 ×10^−11^)^37^ in the *LINC00917–FOXF1* locus is an islet eQTL associated with elevated expression of *FOXF1* (Z-score = 2.12, p = 0.03)^41^ and its neighboring gene *FOXC2* (Z-score = 2.01, p = 0.04)^41^. We identified >20-fold higher *FOXF1* expression in T2D vs. ND β-cells (log_2_FC = 4.38, q = 3.5 ×10^−8^) with no significant difference in *FOXC2* expression (log_2_FC = −0.72, q = 0.74), supporting a causal and concordant role for *FOXF1* induction in T2D genetics and pathophysiology. Similarly, rs67897819 T2D risk allele ‘A’ (OR = 1.07, p = 1.7 ×10^−68^)^37^, upstream of *HNF4A,* is associated with decreased expression of *SGK2,* a serum and glucocorticoid inducible kinase gene 800 kb upstream of *HNF4A* (Z-score = −2.02, p = 0.04)^41^, but not *HNF4A* itself (Z-score = −0.46, p = 0.65)^41^. Consistent with the T2D risk allele effects on *SGK2* expression, it was also reduced in T2D vs. ND β-cells (log_2_FC = − 1.36, q = 8.5 ×10^−4^), which was a stronger difference than *HNF4A* expression (log_2_FC = −1.01, q = 0.04). Expression of neighboring genes (±500kb of rs67897819) did not differ significantly between T2D and ND β-cells (*TOX2, OSER1, GDAP1L1, FITM2, TTPAL, SERINC3, PKIG, ADA* and *KCNK15*), highlighting *SGK2* as a candidate T2D genetic and pathophysiologic causal/effector gene in this region. SGKs, alongside AKT, are activated downstream of mTORC2 (a regulator of β-cell mass) in response to insulin and phosphorylate *bona-fide* AKT target FOXO1^44^, which serves as an anti-apoptotic signal^45^. *SGK2* is a pancreas, liver, and kidney-restricted isoform^46^ that is linked to PD-L1 signaling^47^ and inhibits ferroptosis^48^. Thus, diminished *SGK2* expression in T2D β-cells may contribute to aberrant activation of multiple cell death pathways. This approach also nominated *RASGRP1 and KCNH6* as candidate causal genes whose reduced expression contribute to T2D genetic risk and/or pathophysiology by increasing β-cell susceptibility to pathophysiologic stress and/or enhancing apoptosis propensity. Apoptosis is elevated in *RASGRP1^-/-^* human embryonic stem cell-derived β-cells^49^, and *Kcnh6^-/-^* mice exhibit increased β-cell ER stress, calcium handling defects, and apoptosis that manifests as impaired glucose tolerance^50^. *KCNH6* mutations have been identified in hypoinsulinemic/hyperglycemic patients^51^, and the KCNH6-targeting compound berberine has been shown to stimulate insulin secretion^52^. Mere detection of shared or colocalized eQTL- and T2D-associated variants in non-diabetic islets does not confirm the candidate causal/effector gene’s involvement in mediating T2D predisposition or its functional role. However, our investigation into gene expression differences in T2D β-cells provides complementary evidence supporting roles for these putative T2D causal/effector genes in T2D pathophysiology. Moreover, previous studies for some of the genes identified in these analyses, such as *KCNH6*, support the provocative hypothesis that these genes represent key actors in islet dysfunction with druggable therapeutic potential^50,52^.

Eight DEGs (*FXYD2*, *RPL39L*, *DLK2*, *ITGA1*, *P2RX5, PITPNM2*, *HNF1A-AS1,* and *LTA*; **Figure 3a**, blue) were linked to independent T2D association signals with both concordant and discordant expression effects. For example, rs28413626 T2D risk allele ‘A’ (OR = 1.03, p = 2.1 ×10^−18^)^37^ was associated with higher expression of *PITPNM2* (a membrane-associated phosphatidylinositol transfer protein involved in insulin secretion)^53^ in whole islets (Z-score = 2.1, p = 0.04)^41^, while rs1260294 T2D risk allele ‘T’ (OR = 1.04, p = 2.2 ×10^−15^)^39^, distinct from rs28413626 (LD r^2^ = 0.28, all ancestries combined)^54^ was associated with lower islet *PITPNM2* expression (Z-score = −2.18, p = 0.03)^41^. In T2D β-cells, *PITPNM2* expression was reduced compared to ND β-cells (log_2_FC = −0.72, q = 0.004). Such counteracting signals may reside in distinct regulatory elements within a gene, such as promoters, enhancers or silencers which modulate its expression in diverse ways and lead to opposing effects on gene expression.

We also identified 14 genes, including *GRP, SIX6, SCN3A, ADRB1,* and others, whose altered expression was significantly associated with both T2D genetic and pathophysiologic differences but in opposite directions (**Figure 3a**, gray). For example, rs1895701 T2D risk allele ‘C’ (OR = 1.03, p = 2.4 ×10^−17^)^37^ located between *GALNT3* (upstream) and *CSRNP3*, *SCN2A* and *SCN3A* (downstream) was associated with lower whole islet *GALNT3* expression (Z-score = −3.62, p = 2.9 ×10^−4^)^41^, *SCN3A* (Z-score = −2.77, p = 0.006)^41^ and *SCN2A* (Z-score = −2.46, p = 0.01) ^41^. However, T2D β-cells showed elevated *SCN3A* expression (a sodium voltage-gated channel^22^, log_2_FC = 0.96, q = 3.4 ×10^−5^) and modestly increased *SCN2A* expression (log_2_FC = 0.49, q = 0.04), but no difference in *GALNT3* (log_2_FC = 0.02, q = 0.97) or *CSRNP3* (log_2_FC = 0.07, q = 0.83) expression. This suggests that eQTL associations for such genes exhibiting ‘opposite effects in different tissues’ may be dynamic and context-dependent, potentially switching with the onset of T2D. Indeed, similar context-specific eQTL dynamics have been observed during cell differentiation^55^. Alternatively, this divergent pattern of effects could imply that β-cells, the primary focus of our analysis, may not be the major or exclusive cell type contributing to these eQTL effects. Given that TIGER eQTL associations were detected in whole islet RNA-seq, which represents composite expression from multiple cell types, these genetic variants may exert significant influence in other non-β islet cell types. Opposite eQTL effects between closely related tissues are relatively common^52^. Single-cell dissection of islet cell type-specific regulation (caQTL) and expression (eQTL) should help to resolve these questions and possibilities.

In addition to establishing T2D genetic-pathophysiologic links to identify high priority candidates, we evaluated protein levels of T2D β-cell DEGs from a recent T2D vs. ND human islet proteomic study^56^. We identified 21 genes with concordant differences in both the β-cell mRNA and islet protein levels in T2D vs. ND donors (**Figure 3b**), including a subset with causal links to islet dysfunction and T2D. For example, concordantly T2D-upregulated *SEPT9* gene (and protein) disrupted insulin secretion when overexpressed in rat INS-1(832/13) β-cells^57^, and Cystic Fibrosis transmembrane conductance regulator (CFTR) regulates glucose-dependent electrical activities in β-cells^58^. T2D-downregulated gene/protein *ARG2*, a manganese metalloenzyme, has been linked with polyamine synthesis and regulation of β-cell function^59^, and recent multi-ancestry studies linked rs11114650 T2D risk allele to decreased *LIN7A* expression^60^. Finally, these data support concordant genetic-pathophysiologic effects (**Figure 3a**) implicating reduced *PITPNM2* expression and protein levels in islet β-cell dysfunction and failure. Furthermore, *GRAMD2B*, a downregulated T2D β-cell gene (q = 7.2 ×10^−6^), was also down-regulated at the protein level in T2D islets. GRAMD family proteins are ER-plasma membrane contact site proteins that regulate intracellular Ca^2+^ dynamics^61^. Downregulation of *GRAMD2B* in T2D β-cells likely causes disruption of Ca^2+^ signaling, compromising β-cell function in T2D. Although *GRAMD2B* has not been directly linked with T2D previously, it was identified as a DEG in T2D islets^15^.

To link promising β-cell DEGs to (patho)physiologic effects, we explored diabetes-relevant phenotypes from the International Mouse Phenotyping Consortium (IMPC)^62^, which aims to characterize the function of every protein-coding gene in the mouse genome through whole body gene knock out (KO) mouse lines. 198/511 DEGs identified were tested by IMPC. Homozygous KO of 35 DEGs in mice caused prenatal lethality (n=22) or sub-lethal fitness phenotypes (n=13), highlighting their roles in essential cell survival processes (**Supplementary Table 14**). Importantly, germline deletion of 13 additional DEGs resulted in glycemic defects characteristic of T2D etiology (**Figure 3c, Supplementary Table 14**). To evaluate the effectiveness of mouse KO models in assessing the functional significance of T2D DEGs, we investigated the loss of well-established T2D genes, *Abcc8* and *Kcnj11*, which showed significant glucose tolerance impairment (**Figure 3d)**. Among the identified DEGs, *BNIP3*, a mitochondrial protein originally characterized as an apoptosis inducer^63^, was downregulated in T2D β-cells (q = 0.004, a newly identified DEGs) and *Bnip3^-/-^*mice exhibit impaired glucose tolerance (p = 4 ×10^−9^, **Figure 3d**)^62^ and increased circulating insulin levels (p = 2 ×10^−7^)^62^ compared to wild-type mice. BNIP3 has a dual role in cell fate regulation, balancing between cell death and survival. Notably, during cellular stress (e.g., hypoxia), BNIP3 preserves mitochondrial functional integrity by removing damaged mitochondria via autophagy/mitophagy, thereby protecting cells from death^64^. Loss of *BNIP3* expression in T2D implies that disruption of BNIP3’s stress-responsive adaptation may impair cellular homeostasis. KO of *SLITRK1*, another newly identified T2D upregulated β-cell DEG (q = 0.006) improved glucose tolerance in mice (p = 2.2 ×10^−6^, **Figure 3d**)^62^. *SLITRK1* is a neuronal transmembrane protein linked to neurological diseases such as schizophrenia^65^ and Tourette’s syndrome^66^ and promotes excitatory synapse formation when overexpressed in cultured rat hippocampal neurons^67^. Its upregulation in T2D connects β-cell dysfunction to an altered islet-nerve communication which is critical in fine-tuning insulin secretion in response to circulating glucose levels. GRAMD2B, a downregulated mRNA/protein in human T2D β-cells/islets is crucial for glucose hemostasis, as *Gramd2b^-/-^* mice exhibit impaired glucose tolerance (p = 2.3 ×10^−5^, **Figure 3d**)^62^. Thus, our integration of mouse physiologic data provided insights into *in vivo* gene functions and systemic (patho)physiologic effects of identified DEGs, enabling identification of previously unrecognized genes with crucial roles in T2D physiology as promising therapeutic targets.

### T2D-associated differences in putative islet β-cell subpopulations

Single-cell and targeted islet analyses have reinvigorated the interests in and debates around models in which (patho)physiologic states such as T2D are characterized by molecularly and functionally distinct endocrine cell subpopulations or states with variable maturity, stress, hormone secretion and glucose responsiveness^68,69,70,71,72^. Thus, we sought to identify robust, reproducible endocrine cell subpopulations present in our 48-donor cohort and assess if they are significantly altered in T2D, PD, or ND states. We analyzed 74,812 α-, 99,029 β-, and 10,770 δ-cells from T2D, PD and ND islets and identified seven putative α- and δ-cell subpopulations and eight β-cell subpopulations (**Supplementary Table 15** and **Figure 4a, Supplementary Figures 8a-d** and **9a-d**). All β-cell subpopulations expressed *INS* at comparable levels confirming their β-cell identity, with no variation in clustering between sexes, ancestries or sequencing chemistries (**Supplementary Figure 10a-c)**. Each cell type subpopulation was detected in every donor across the three glycemic states (**Figure 4b** and **Supplementary Table 15**). Within each pancreatic islet endocrine cell type, we observed subpopulations with endoplasmic reticulum (ER) stress and/or hypoxia (a critical ER stressor) response gene expression signatures: clusters 7 and 2 (α), clusters 6 and 2 (β) and, cluster 4 (δ) (**Supplementary Table 15**, **Figure 4c, Supplementary Figures 8e and 9e**), implying a broad role of ER homeostasis perturbations or cycles in multiple islet cell types^73^, not just β-cells. Indeed, ER stressed β-cells with elevated *DDIT3, HSPA5, HERPUD1,* and *TRIB3* expression have been previously documented (**Supplementary Figure 10d**)^74,75,76^. Additionally, we detected a putative β-cell subpopulation with high functional capacity and/or output exhibiting elevated expression of genes enriched in the ‘insulin secretion’ pathway that harbor a spectrum of diabetes-associated sequence variation – *ABCC8, G6PC2, PDX1, SLC30A8, RBP4* (cluster 1). We also identified β-cell clusters with expression signatures associated with translation initiation (cluster 3); heat shock proteins (cluster 4, proliferative vs. mature cells with elevated *CFAP126* (*Fltp* in mice) expression^77^); regulation of signaling receptor activity (cluster 5, CD63^hi^ cells with enhanced glucose-stimulated insulin secretion^78^); cellular senescence (e.g., *CDKN2A*, *CDKN2B*, *PLK2*, *B2M* expression, cluster 7); and cellular transport (cluster 8, also found in α and δ cells) (**Supplementary Table 15**).

**Figure 4:**
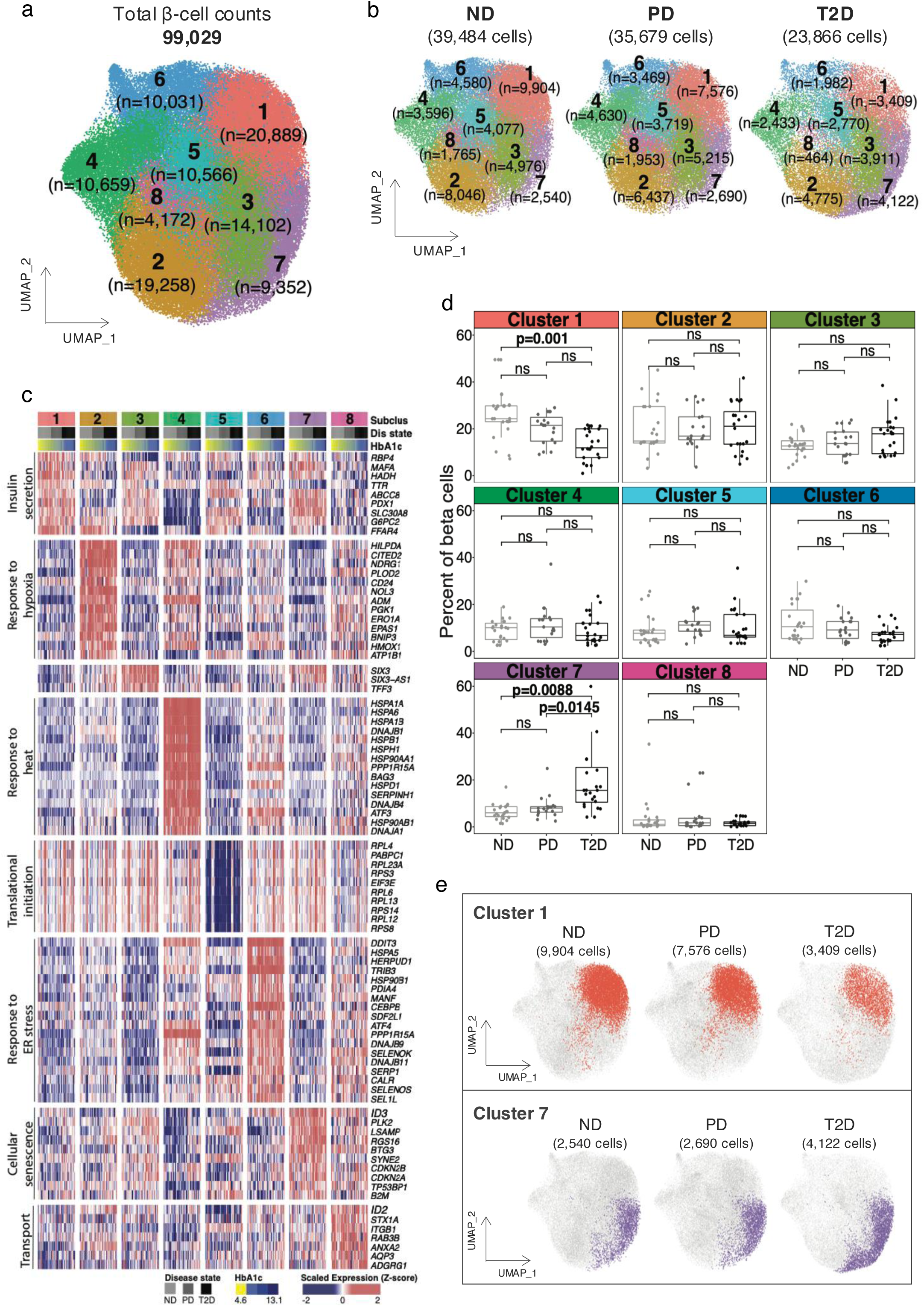
β-cell subpopulation differences in T2D vs. ND and PD islets. **(a)** Sub-clustering analysis of 99,029 β-cell transcriptomes reveals eight putative subpopulations. n=number of cells in each sub-population. **(b)** Dot-and-box plots showing β-cell sub-population distributions in ND, PD, and T2D samples. **(c)** Heatmap of scaled marker gene expression (rows; y-axis) for each β-subcluster (x-axis) aggregated by donors (columns). For each subpopulation, donor profiles are grouped into ND (light gray), PD (dark gray), and T2D (black) states, and then sorted based on ascending HbA1c levels. Enriched biological processes (left) associated with representative differentially expressed genes (right) are shown for each subpopulation. **(d)** Dot-and-box plots comparing per-donor percentages of each β-cell subpopulation in ND, PD, or T2D individuals. Each dot represents a donor. Bonferroni-adjusted p values from Tukey’s honestly significant difference test are reported for significant differences; ns=not significant. **(e)** UMAP plots illustrating inversely correlated changes in cluster 1 (decreasing) and cluster 7 (increasing) cells within the bulk β-cell cluster from ND to PD to T2D states.

α- and δ-cell sub-clusters showed no significant difference between T2D, PD, or ND donors (**Supplementary Figures 8f** and **9f**). Although no β-cell subpopulations were uniquely present or absent in T2D donor islets, two exhibited significant reciprocal quantitative differences in their relative abundance between T2D vs. ND or PD donors (**Figure 4d and 4e**). The putative high-functioning, ‘insulin-secreting’ cluster 1 population proportions were reduced an average of 10.5% in T2D vs. ND β-cells (p = 0.001), underscoring the potential importance of this sub-population for proper islet function and/or its enhanced sensitivity to pathophysiologic changes. In contrast, the proportion of ‘cellular senescence’ cluster 7 cells increased by an average of 12.3% in T2D vs ND β-cells (p = 0.009); this significant increase was also observed in T2D vs. PD β-cells (p = 0.02, average increase = 9.7%). These unsupervised subpopulation analyses thus support emerging reports of increased β-cell senescence in T2D^79,80,81^. Together, these subpopulation shifts combine with 10-15% overall reductions in T2D donor β-cell numbers/proportions in this cohort (**Figure 1e**) to result in ∼25-30% reduction of functional β-cell mass.

## Discussion

Here, we report comprehensive profiling and comparative analyses of approximately a quarter million human islet single-cell transcriptomes from a unique, HPAP-independent 48-donor cohort representing ND, PD, and T2D states. These studies revealed significant T2D-associated changes in β-cell (sub)populations that collectively manifest as ∼25-30% reductions in functional β-cell mass compared to ND or PD islet donors. Moreover, we identify 511 genes whose expression is perturbed in T2D β-cells. T2D genetic and pathophysiologic factors exert convergent and concordant effects on their expression for 58 of these T2D β-cell DEGs, nominating them as high priority candidate causal and putative interventional target genes. We provide these human islet single cell transcriptomic data as an accessible resource to the islet biology and diabetes research communities in two formats – the free-to-use CellxGene (https://cellxgene.cziscience.com/collections/58e85c2f-d52e-4c19-8393-b854b84d516e) platform for investigating and retrieving gene expression signatures across various islet cell types in the context of T2D and the Transcriptome Atlas of Pancreatic Islet Cells (TAPIC; https://thejacksonlaboratory.shinyapps.io/TAPIC_Stitzel_Lab/) R shiny app we created for data visualization – to enhance exploration and integration of this comprehensive and disease-relevant islet transcriptome atlas.

This study significantly advances ongoing efforts aimed at important and widely editorialized gaps in our collective ability to identify reproducible “T2D genes” in human islets. We applied a pseudobulk analysis approach to identify 746 T2D β-cell DEGs at FDR<5%, 511 of which differed >1.5-fold in expression. One-third of these 511 robust T2D DEGs have been reported in previous studies, including those such as *DGKB, ASCL2, GOLT1A, ARG2, PPP1R1A*, and others reported as islet and/or β-cell T2D DEGs in HPAP and independent cohort studies. Thus, continued efforts and increasing sample sizes are homing in on an ever-increasing set of high-confidence T2D-associated DEGs as anticipated. Systematic efforts to collect and uniformly process, QC, and analyze all human islet single cell profiles, such as those of the newly established Pancreas Knowledgebase (PanKBase), should build on this momentum and enhance our multi-omic understanding of islet cell type-specific (dys)function.

Neuroreceptor signaling and vitamin A metabolism emerged among the most significant up- and down-regulated pathways/processes among DEGs, suggesting substantive alterations in both neuroactive ligand receptor signaling and metabolism in T2D β-cells. Although both have been linked to islet development and function by targeted studies in model systems, the gene-based enrichments detected here uniquely highlight their empiric links to human islet pathology and T2D. Cnop and Pipeleers noted two decades ago the protective effects of vitamin A vs. LDL-induced toxicity in rat islet β-cells^82^. Chemical screens and mechanistic studies in zebrafish added retinoic acid biosynthesis and signaling alongside Notch as important contributors to islet differentiation and β-cell regeneration^83–85^. Subsequent rodent studies demonstrated that vitamin A deficiency increased β-cell apoptosis, increased α−cell mass and hyperglucagonemia^86^, while dominant negative-negative RAR-α inhibition led to age-dependent decreases in plasma insulin resulting from impaired GSIS, decreased β-cell mass and per-β-cell insulin content^87^.

Expression of genes encoding several neurotransmitter receptors - adrenergic (*ADRB1*), glutaminergic (*GRIN1*, *GRIN2A*), cholinergic (*CHRNA3*, *CHRNA5*), purinergic (*P2RY1*, *P2RX5*), serotonin (*SSTR1*), GABAergic (*GABRA2*), and glycine (*GLRA1*, *GLRA3*) - were altered in T2D β-cells. β-cell function is tightly controlled by the autonomic nervous system^88,89^ and neurotransmitters, whether synthesized locally within the islet or released by the neurons, are crucial for regulating insulin secretion. Glutamate, for instance, is an excitatory neurotransmitter that signals through N-methyl-D-aspartate receptors (NMDARs), which are ligand-gated cation channels with high calcium permeability encoded by *GRIN* genes^22^. β-cell-specific deletion of *Grin1* reveal enhanced GSIS and improved glucose tolerance in mutant mice^90^. In humans, treatment with the NMDAR antagonist dextromethorphan (DXM) increases serum insulin levels and improves glucose tolerance in individuals with T2D^90^; pancreatic NMDARs are therefore being evaluated as promising therapeutic targets for diabetes management. Furthermore, gamma-aminobutyric acid (GABA) is simultaneously released with insulin by β-cells and binds β-cells GABAA receptors to inhibit insulin secretion as an autocrine signal^91^. *GABRA2* encodes the exclusive GABAA receptor of human β-cells^92^ and has been reported to feature relatively closed chromatin conformation in T2D islets compared to non-diabetic islets^93^. Neurotransmitter receptor dysfunction in T2D islets has been discussed before^94^. Given their critical involvement in β-cell signaling and insulin secretion, these neuroreceptor genes, which are currently under-studied in the context of diabetes, warrant further exploration.

Using a combination of genetic and comparative analyses along with experimental approaches, we nominated 92 T2D β-cell DEGs as prioritized candidate causal T2D genes (44 reported and 48 newly identified). This prioritized list includes genes with long-standing gain- vs. loss-of-function T2D GWAS variant effects on islet gene expression (*DGKB, ST6GAL1 vs. STARD10*) and exciting new candidates that span the variant-to-function gamut like *KCNH6* and capture concordant multi-study, multi-modal effects, such as *GRAMD2B.* Several T2D β-cell DEGs exhibit diabetes-relevant glucose homeostasis phenotypes in knockout mice that are directionally consistent with their dysregulation in T2D vs. ND islets, such that germline knockout of up- or down-regulated genes improve (e.g. *SLITRK1, IRF8*) or impair (e.g., *GRAMD2B, BNIP3*) glucose homeostasis, respectively. Systematic and targeted mechanistic studies of these prioritized genes in primary human islets and using β-cell-specific knockouts are warranted and will firmly establish their causal roles in islet dysfunction and T2D.

Although we identified robust and extensive gene expression changes in T2D β-cells, we detected surprisingly minimal-to-negligible alterations in T2D islet α− or δ-cells. Our comparative analyses and experimental validation support these T2D β-cell DEGs as compelling causal candidates. Although the biological vs. potential technical basis for the surprising lack of differences in the other cell types is unclear, it is noteworthy that this apparent conundrum was also observed in recent independent, HPAP cohort-based analyses^6^. All donor islets in this study were handled and processed using standardized protocols after overnight recovery from shipping and islet cell types were captured in parallel with comparable sequencing depth, so it is unlikely this discrepancy results from capture or sequencing bias in the platform. Despite best efforts to match donor characteristics, these experiments and analyses involve *ex vivo* islets from donors with inherently variable lifestyles and environments. It is possible that 5.5 mM glucose concentrations in standard culture conditions may provide stimulation or stress that “unmasks” the β-cell differences and deficiencies observed but not those in other cell types. For example, α-cells respond to low (0mM) glucose and amino acids, so islets may need to be cultured in these conditions to “unmask” α-cell deficiencies in T2D islets. Future studies comparing T2D vs. ND islet cell differences with cell type-specific stimuli/stressors will be important to test this hypothesis and could enhance our understanding of non-β-cell contributions to islet dysfunction in T2D.

Unsupervised analyses of single cell transcriptomes for each islet cell type identified multiple putative β-cell subpopulations, present in every donor of this cohort, including several previously described^72,74–78,95,96^. We confirm and extend to α− or δ-cells the widely reported ER stress signature β-cell sub-populations described in multiple reports^97–99^, perhaps reflecting more generalizable cycles of hormone production and secretion, and identify reported subpopulations with elevated *CD63*^78^ and *CFAP126 (*the human *Fltp* orthologue)^77^ expression. However, the abundance/proportions of these subpopulations did not differ between T2D and ND or PD individuals, suggesting that they may contribute to physiologic rather than pathologic β-cell heterogeneity. In contrast, we detected significant T2D-associated increases in β-cells exhibiting an elevated senescence signature gene expression signature. Senescence has been recently implicated as a (mal)adaptive β-cell process in both type 1^100^ and type 2 diabetes^101,102^; changes in this putative subpopulation may underlie reported intra-individual heterogeneity in *CDKN2A* expression and nearby chromatin accessibility in T2D β-cells from recent trajectory-based analyses of a smaller cohort^8^. Mechanistic studies of this putative T2D-associated β-cell subpopulation and the effects of senolytic vs. senomorphic agents on T2D islet function are warranted to determine and discern their helpful vs. harmful role(s). More broadly, spatial, *in situ-*based approaches will be critical to assess and compare within- vs. between-islet organization of these putative subpopulations, and their effects on islet function, in distinct regions of the pancreas (e.g., head, neck, body, tail) or healthy vs. diabetic individuals.

## Funding

This study was made possible by generous funding from the American Diabetes Association Pathway to Stop Diabetes Accelerator Award (1-18-ACE-015) and National Institutes of Health (NIH) award number R01DK118011 (to M.L.S) as well as Department of Defense Congressionally Directed Medical Research Program (CDMRP) award number W81XWH-18-0401 (to M.L.S. and D.U.). C.N.S. was also supported by American Diabetes Association grant 11-22-JDFPM-06. Opinions, interpretations, conclusions, and recommendations are solely the responsibility of the authors and do not necessarily represent the official views of ADA, NIH, or DOD. We gratefully acknowledge contributions of JAX Single Cell Biology and Genome Technologies services and Research Cyberinfrastructure computational resources at The Jackson Laboratory for expert assistance with the work described in this publication. We are indebted to the anonymous islet organ donors and their family, which were provided by the NIDDK-funded Integrated Islet Distribution Program (IIDP) (RRID:SCR_014387) at City of Hope (2UC4DK098085). Special thanks to Dr. Raphael Scharfmann at Institute Cochin for help optimizing EndoC-βH3 culture. We thank Ucar and Stitzel lab members for critical feedback throughout this study.

## Data and Code Availability

All human islet sample and single cell RNA-seq datasets have been deposited in the BioProject and Gene Expression Omnibus databases under accession numbers PRJNA913127 and GSE221156. The data have been processed in the R statistical package and the analytical pipeline including the detailed methodology, the code and the associated plots/tables are available at Zenodo under https://zenodo.org/records/14656366. The reader can use our code outlined in the Pipeline_html.Rmd and Pipeline_html.html files to replicate the results and to explore other aspects of our data discussed in detail in the manuscript.

At the single-cell level, the processed data are available for interactive visualization by cellxgene at https://cellxgene.cziscience.com/collections/58e85c2f-d52e-4c19-8393-b854b84d516e. The dataset is divided into four instances, one referring to the data of all annotated cells and one for each of the major identified cell types, namely Beta, Alpha and Delta cells. At the pseudobulk level, processed data are available for interactive visualization by the TAPIC Rshiny applet at https://thejacksonlaboratory.shinyapps.io/TAPIC_Stitzel_Lab/.

## Conflict of interests

The authors declare no competing interests.

## Methods

### Single cell library preparation and sequencing

Pancreatic islets from 48 individuals consisting of 17 ND, 17 T2D and 14 PD donors were cultured using CMRL, supplemented with 10% FBS, 1% Glutamax,1% Pen/Strep for 14 days. Islet-derived fibroblasts were harvested and gDNA extracted using the Blood & Tissue kit (Qiagen). The RNAse A (Qiagen) treated genomic DNA samples were genotyped using the Infinium Global Diversity Array-8 v1.0 Kit (Illumina). Single cell capture, barcoding, and library preparation were performed using the 10X Chromium platform (https://www.10xgenomics.com) according to the manufacturer’s protocol for chemistries v2 (#CG00052) and v3 (#CG000183). Illumina base call files for all libraries were converted to FASTQs using CellRanger-6.1.2 demultiplexing and count pipelines (https://www.10xgenomics.com). Initially, we used cellranger’s *mkfastq* to demultiplex the raw base call (BCL) files generated by Illumina sequencers, perform adapter trimming, and retrieve the 10-bp length UMI bases to be included into the generated FASTQ files for downstream processing. We processed that raw FASTQs with STARsolo^103^ using STAR 2.7.9a (https://github.com/alexdobin/STAR/blob/master/docs/STARsolo.md). The barcode demultiplexing was done with the default V2 / V3 whitelists coming from the CellRanger v.6 installation (https://kb.10xgenomics.com/hc/en-us/articles/115004506263-What-is-a-barcode-whitelist-). For each of the Gel bead-in Emulsions (GEMs), we aligned the reads to the full Ensembl human genome GRCh38 (https://uswest.ensembl.org/Homo_sapiens/Info/Index) and used the standard STAR spliced read alignment algorithm to assigned them into the exonic, intronic and intergenic groups. We performed error-correction and deduplication of the Unique Molecular Identifiers (UMIs) and quantified the per-cell gene expression to generate the raw UMI data for each library. To filter out the empty droplets we employed the EmptyDrops_CR background model^104^ that generated the filtered UMI data of *G* = 36,601 genes (both protein-coding and non-coding) and *N*\* = 414,082 cells across the *J* = 54 libraries for downstream processing. The median number of cells across libraries was 7,748 with a 25%-75% Inter-Quantile Range (IQR) of 5,891-9,133. Our data exhibited the typical high-quality features suggested by the 10x’s guidelines (CG000329-Rev A, Technical Note): fraction of reads with valid barcodes (ideal: >0.75; our median: 0.98), fraction of UMI bases with Q-score ≥30 (ideal: > 0.65; our median: 0.96) and fraction of unique reads in cells (ideal: > 0.70; our median: 0.81).

### Experimental design and meta data information

The similarity of the three glycemic states was assessed in terms of the Euclidian dot product 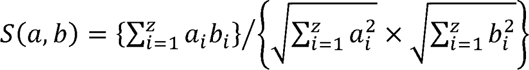, where *a, b* are the z-dimensional attribute vectors of a glycemic state pair under comparison (z = 4) and *a_i_, b_i_* are the ith components of these vectors (one of sex, ancestry, age and BMI). We generated *J* = 54 10X libraries containing the data of either single or multiplexed islets. The first 12 islets (in processing date), coming from 4 ND, 1 PD and 7 T2D donors, were generated with the V2 chemistry. Cells from 12 islets of V3 chemistry had their RNA sequenced across multiple islet-specific or genetically multiplexed islet libraries^105^.

### Ambient RNA Decontamination by SoupX

Ambient RNA is the pool of mRNA molecules released in the cell suspension likely from stressed or apoptotic cells. It is incorporated into the droplets resulting in cross-contamination of transcripts between different cell populations. We estimated and removed contamination in individual cells by the SoupX model^106^ using the *soupX-1.6.2* R package from CRAN. We processed each library *j* = 1,…,54 separately. First, we converted STARsolo’s raw and filtered UMI data into respective Seurat v.4 objects^107^ (STAR-Methods) that were subsequently merged into a single SoupX object. Seurat’s filtered data were normalized, scaled and clustered with Leuven on the UMAP reduced representation (see “*Seurat clustering by library*”). The clustering was fed into the SoupX object for ambient RNA contamination estimation and adjustment.

For decontamination, we considered as empty the droplets with less than 10 UMIs^106^ and estimated the fraction of background expression from each gene *g* of library *j* (*j* = 1,…,54) across all empty droplets *E* as 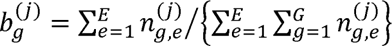 where *n_g,e_* denotes the observed counts for gene *g* in the empty droplet *e*. We used the background to estimate likewise each cell’s *c* contamination fraction as 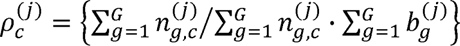 where the sums are taken across all genes *G* in each cell *c* of library *j* (SoupX’s *autoEstCont* method). Finally, the endogenous (decontaminated) counts were retrieved as 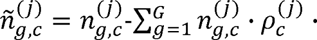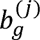. The decontaminated counts of all genes *G* in cells of library *j* were stored in Seurat objects and were used for the downstream analysis. To assess the degree of contamination across all cells *N*\*, we estimated for each gene *g*, *g* = 1,…,36,601 in library *j*, *j* = 1,…,54, the difference in UMI counts before and after decontamination as 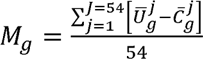, where 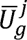 was the average uncorrected UMI of gene *g* across all cells of library *j* and 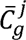 was the average corrected UMI of gene *g* across all cells of library *j*. The *M_g_* levels were examined as a function of the average expression quantity 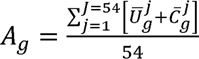 by a classical MA plot that indicated at each *A_g_* level the degree of average decontamination for gene *g* across all libraries. In addition, we estimated the average contamination ranking for each *g* as 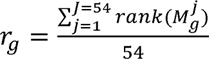, where 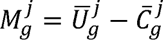. Combining the information, our data separated three broad clusters of contaminants, the most prominent of which included famous endocrine and exocrine markers such as Insulin (*INS*), Glucagon (*GCG*), Somatostatin (*SST*), Pancreatic Polypeptide (*PPY*), Regenerating Family Member 1 Alpha (*REG1A*), Serine Protease 2 (*PRSS2*) and other genes such as Transthyretin (*TTR*) and Islet Amyloid Polypeptide (*IAPP*). The genes of this cluster were ranked on average across all cells among the top 10 contaminants. A second cluster consisted of several mitochondrial and ribosomal genes ranked on average among the top 20 to 100 contaminants.

### Sample deconvolution by Demuxlet

We utilized modern barcoding technology to improve the throughput of detected cells and genes via genetic multiplexing (see “Genetic multiplexing”) for a limited set of 12 libraries generated under the V3 chemistry. Each library consisted of multiplexed barcoded cells from two islets of donors with different clinical and demographic background and processed with Demuxlet^108^. Demuxlet considered the islet’s genetic variation to determine the genetic identity of each droplet through a set of single nucleotide polymorphisms (SNPs). The islet SNPs were identified from the islet genotypes after extended quality control. The sample IDs of the processed genotypes were validated by comparing them to paired bulk ATAC-seq data using verifyBamID (https://github.com/statgen/verifyBamID/releases). To obtain the SNPs, we used plink v1.90^109^ and generated the Extended variant information files (.bim) accompanying the binary genotype information for each chromosome of the GRCh37 human genome. We performed error-correction for each file with HRC-1000G-check-bim (https://www.well.ox.ac.uk/~wrayner/tools/) (STAR-Methods) to remove duplicate variants, mismatched variants, palindromic variants with frequency > 0.4 and to correct strand flips using the reference file PASS.Variantsbravo-dbsnp-all.tab containing 170M variants on ∼15k individuals from the dbSNP database^110^. We joined and sorted the error-free data of all chromosomes with bcftools-1.11 (https://samtools.github.io/bcftools/bcftools.html) and, at the last step, we used liftOver to convert the genotype coordinates in the .vcf file to GRCh38 and obtained the barcode-to-islet associations.

Demuxlet quantified the likelihood that the *C*^(j)^-th droplet of library *j* originated from the *m*_1_ or the *m*_2_ islets that have been multiplexed with mixing proportions (*1*-a):*a*. The likelihood had the form:

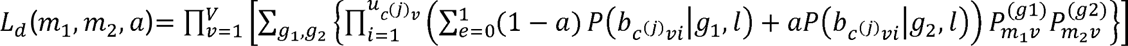

The above expression considers the reads from *C*^(j)^ barcoded cell-containing droplets of library *j* multiplexed across two islets *m*_1_ and *m*_2_. The islet genotypes are available across *V* exonic variants, *u_c_*^(j)^_v_ is the number of unique reads overlapping the *v^th^* variant of the *C*^(j)^ droplet, *b_c_*^(j)^_v_ is the variant overlapping base call from the *i*^th^ unique read, *i* = 1,…,*u*_c_^(j)^_v_ representaing reference (R), alternative (A) and other (O) alleles and *l* is a latent variable indicating whether the base call is correct (0) or not (1).

We used directly Demuxlet’s .best output files that summarized the best assignment of the *C*^(j)^ -th droplet of *j* between two multiplexed islets *m*_1_ and *m*_2_. The provided information explicitly associating each *C*^(j)^ either to a single islet (singlets) or to both (Demuxlet doublet) with high probability. We integrated this information of each library with the Seurat soupX corrected objects and filtered out the Demuxlet doublets from further analysis. Across the 12 libraries, we removed on average 1,065 doublets (25%-75% IQR: 650-1319). We found that the number of Demuxlet doublets was weakly anti-correlated to the library processing date (Pearson’s *rho* = −0.57, p-value = 0.053) and also correlated to the number of STARsolo cell-containing droplets (Pearson’s *rho* = 0.611, p-value = 0.035). We kept 401,305 cells for further analysis across *j*\* = 1,…,66 libraries holding the demultiplexed data of *D* = 48 islets (some islets are represented more than once).

### Quality control by library

The quality control (QC) analysis was performed iteratively on the decontaminated raw counts of each *j*\* = 1,…,66 demultiplexed library. It consisted of the following steps:

1. Preliminary filtering: We filtered out all cells with *nFEA* ≤ 500 or *nUMI* ≤ 1000 or *pMT* ≥ 50% as a first-pass data cleaning for the subsequent pre-processing steps. The identification of high-quality cells combined multi-step doublet estimation, stricter *nFEA*, *nUMI*, and *pMT* cutoffs and statistical testing as shown below.
2. Doublet estimation: We used Scrublet^111^ and DoubletFinder^112^ to estimate the neotypic doublet from each *j*\* library. Scrublet was run in Python 3.6.15 on the Seurat-to-10x formatted data (function write10xCounts of DropletUtils R package) with an expected 10% doublet ratio. We visualized the doublet scores of the observed and simulated doublets in a histogram and inspected their bimodal distributions to set an appropriate cutoff that separates the doublets from the singlets. DoubletFinder operated on the normalized and clustered Seurat data of each *j*\*. Similar to Scrublet, it simulated doublets with a set of user-defined parameters: *p_N_* (proportion of generated artificial doublets), *p_k_* (the PC neighborhood size to compute each cell’s proportion of artificial k nearest neighbors on a PCA, *p_ANN_*), *nPC* (the number of principal components (PCs)) and *nExp (*a threshold to make the singlet / doublet prediction, similar to the expected doublet ratio).
3. Clustering: We followed Seurat’s v4.0 standard pre-processing workflow^108^ from data normalization to cell clustering (see *“Seurat Clustering by library”*) to filter out low quality cells before the main data integration step. We visualized the distribution of *nFEA*, *nUMI* and *pMT* of all cells across each cluster *x* of *j*\* to determine the appropriate cutoffs.
4. Marker analysis: To avoid over-filtering, we utilized marker expression analysis and pre-annotated the clusters with known endocrine and exocrine marker genes (see “Cell Annotation”), considering that *nFEA*, *nUMI* and *pMT* may vary substantially across the various cell types.
5. *pMT* comparisons: We examined whether certain cell types and, more importantly, glycemic states (ND, PD, T2D) exhibited higher *pMT* rates than others to adjust the cutoffs. The comparisons were performed across annotated clusters within each state and across states within each cell type using ANOVA and Tukey’s Honestly Significant Differences (HSD) pairwise tests with Bonferroni corrected p-values.

Scrublet set an automatic threshold at the point between the two modes of the simulate scores. We visually determined that the optimal doublet threshold was at 0.25 in all libraries. For DoubletFinder, we normalized and clustered the Seurat data (10 PCs, Leuven clustering with resolution = 0.5 on the UMAP) and instructed the algorithm to simulate doublets using *p_N_* = 0.25 (default), *p_K_* = 0.09 (estimated), *nExp* = 0.1 and *nPC* = 10 (default). Marker analysis showed that, in contrast to Scrublet, DoubletFinder often assigned higher doublet rates in *PPY*^+^, *PECAM*^+^ and *COL*1*A*1^+^ cell clusters. To avoid over-filtering, we removed only the common Scrublet-DoubletFinder doublets leaving a total of 336,692 cells for further processing (median = 6,663; 25%-75% IQR = 4,965 –8,288).

### Seurat clustering by library

The raw counts of each library were normalized with the *LogNormalize* model that employed a global-scaling (library size) normalization with *scale.factor* = 10,000 and log-transformed the result. The top 2,000 variable features, exhibiting the highest cell-to-cell variation, were extracted and their normalized counts were scaled to fit on a Principal Components Analysis (PCA) model for linear dimensionality reduction. We used standard quality control on the PC loadings to empirically determine the optimal number of PCs accounting for the data variability. For each library, we visualized the PC loadings and the associated heatmaps of gene expression and kept the first 100 PCs for the UMAP representation. We constructed a shared nearest neighbor graph by calculating the neighborhood overlap (Jaccard index) between every cell and its 20 nearest neighbors obtained from the cell Euclidean distances. We clustered the data with Leuven and clustering resolution parameter equal to 1. We inspected separate violin and 2d plots of *nFEA* vs *pMT* and *nUMI* vs *pMT*. Based on the spatial 1d and 2d patterns, we flagged all cells with *nFEA* < 1400 unless, at these cutoffs, the flagged cells enriched for an annotated cluster (high-quality exocrine cell types had relatively low number of features) in which case, we determined that *nFEA* < 1000 was the most appropriate cutoff.

To set the *pMT* cutoff, we compared various quantiles of the *pMT* distribution across annotated cell types (derived by merging the cells of the above annotated clusters) and glycemic states (ND, PD, T2D). We used ANOVA and Tukey HSD pairwise tests to find whether the *pMT* rates differed significantly across the cell types. The analysis showed that α- and β-cells exhibited statistically higher *pMT* medians, 70-th quantiles and 90-th quantiles compared to other types (see “*Quality control by library”*) at Bonferroni adjusted p-value 5%. On the other hand, none of the cell types showed significant differences among the *nUMI*s of ND vs PD vs T2D at Bonferroni adjusted p-value 5%. Based on these plots and results, we flagged all cells with *nUMI* > 40%. All flagged cells were removed from further analysis reducing the dataset to 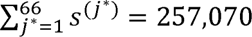 cells (median: 5,135; 25%-75% IQR: 4,063-6,014).

### Data Integration and final cell filtering

We used Harmony v3.8^113^ within the Seurat v4.0 workflow to integrate the *j\** = 1,…,66 libraries each consisting of *c^j*^* cells (after quality control by library). Harmony exhibits excellent scaling properties for large populations and accommodates complex experimental designs allowing the user to explicitly specify the model parameters (factors) to be integrated. First, we merged the normalized data across all 66 libraries (see *“Seurat Clustering by library”*) and extracted the top 2,000 variable features for scaling. The cells were embedded in a 100-dimensional PCA space and Harmony adjusted iteratively for sex, chemistry, and ancestry until convergence (10 iterations). At each iteration, the method used fuzzy clustering to assign each cell to multiple clusters while preserving the data diversity via the Ω(.) term that penalized statistical dependence between batch identity and cluster assignment. The estimated cluster and batch centroids were used to derive a batch correction factor and a cell-specific correction factor which were iteratively updated until convergence to a stable clustering representation. The procedure generated 36 cell clusters. The median ratio of females across the 36 clusters was 0.36 (25%-75% IQR: 0.3 – 0.41) which was close to the overall female: male ratio of 0.37:0.63. Similarly, the median values were 0.81 for V3 chemistry (overall: 0.79), 0.49 for the European ancestry group (overall: 0.47), and 0.34 for the Hispanic ancestry group (median: 0.37).

### Doublet enrichment

We estimated which of the Harmony integrated clusters were enriched in Scrublet doublets by the Fisher test. For each integrated cluster *x*\*, *x*\* = 0,1,…,35, we calculated the doublet ratios 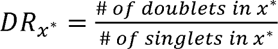 and 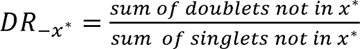. The enrichment test was performed on the 2 *x* 2 confusion matrix with column entries the numerator and denominator of each of the above quantities. The clusters with *OR* = *DR_x*_*:*DR_-x*_* > 2 and Fisher test *FDR* ≤ 1% were those labeled as significantly enriched. Clusters 29 with *DR*_29_ = 1.248 and 20 with *DR*_29_ = 0.307 stood out with relatively high ratios. The Fisher enrichment test showed that clusters 29, 20, 15, and 23 were significantly enriched in doublets (odds ratio, *OR* > 2; Fisher test FDR<1%). We ran differential expression analysis at the single-cell level with Seurat’s logistic regression model comparing the average expression of gene *g* in cluster *x*\* against its average expression in all other clusters after adjusting for sex, chemistry and ancestry (*logFC* ≥ 0.25 and *FDR* ≥ 1%). Combined with evidence from marker expression analysis, we found that cluster 29 expressed highly both *INS* and *SST* while cluster 20 expressed both *GCG* and *SST*. None of the other clusters showed such evidence. We labeled all cells of 20 and 29 as doublets and filtered them out along with all other Scrublet doublets, ending up with the final set of *N* = 245,878 high-quality cells for downstream processing. We summarized the number of cells after each filtering step by islet and examined whether our strategy and cutoffs led to differences in cell percentages of each step across the glycemic states. We used ANOVA and Tukey’s HSD to test the null hypothesis 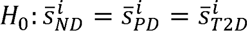 versus the alternative that at least one of the 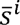 differed, where 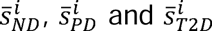 were the average percentage of cells across the ND, PD and T2D islets, respectively, after filtering step *i*. We did not detect any statistically differences at Bonferroni adjusted p-value = 5%.

### Doublet estimation

We estimated three types of doublets from our data. First, for the genetically multiplexed libraries, we found Demuxlet doublets consisting of cells from different donors (see “*Sample deconvolution by Demuxlet”)*. Second, within each library, we utilized Scrublet and DoubletFinder that simulated doublets from the raw UMI counts (after Demuxlet when applicable) and, via nearest neighboring clustering, calculated the likelihood for an observed cell to be a doublet (“Quality control by library”). Third, after data integration, we identified clusters enriched in Scrublet doublets and expressing highly more than one of the known endocrine and exocrine markers. All cells of such clusters were potential doublets (*“Doublet enrichment”*).

### Cell Annotation

We annotated the integrated UMAP clusters of high-quality cells with a large list of known endocrine and exocrine marker genes obtained from the literature and by differential expression analysis. The latter was conducted by Seurat’s logistic regression model that compared gene’s *g* average expression in *x*\* against its average expression all other clusters after adjusting for sex, chemistry and ancestry. We reported significant genes at *logFC* ≥ 0.25 and *FDR* ≥ 1%. The estimated cell types (and indicative markers) were beta (*INS*), alpha (*GCG*), delta (*SST*), gamma (*PPY*), epsilon (*GHRL*), ductal (*KRT19*), acinar (*REG1B*), stellate (*COL1A1*), activated stellate (*FABP4*), endothelial (*PLVAP*), Schwann (*NGFR*), immune (*C1QC*), mast (*TPSB2*) and proliferating cells (*TOP2A*). For each islet, we calculated the number of cells across the glycemic states of each cell type. We tested whether the absolute frequencies and/or the cell percentages differed significantly in ND vs PD vs T2D by ANOVA and Tukey’s HSD pairwise tests with Bonferroni correction.

### Conversion to pseudo-bulk

We considered that the single cells within an islet are not independent of each other and estimated the transcriptomic differences across cell types and glycemic states at the pseudo-bulk level where the islets served as the biological replicates. We aggregated the single-cell raw counts and meta data (clinical and demographic information) associated with each cell type *z* and islet *p*, *p*= 1,…, *P*, and generated *Z* = 14 pseudobulk RNA-seq count matrices of dimension *G* x *P_Z_*, where *P_Z_* might differ across the cell types. The aggregated data were obtained by the *aggregate.Matrix* function of the *Matrix.util* v0.9.8 R package. The quality control of the pseudobulk data was done in terms of the islet library sizes (*nUMI*), the islet number of detected genes (*nFEA*), the percentage of reads mapped to the MT genome (*pMT*) and the Counts-per-Million (CPM) normalized log2-expression profiles of the top 50 most expressed genes. We employed 2d Multidimensional Scaling (MDS) in edgeR v3.34.1^114^ to detect the major sources of variability within each cell type.

### Differential Expression Analysis with edgeR

We quantified the transcriptomic differences of the cell types and the glycemic states of each endocrine cell type by edgeR on the pseudobulk level. Each model adjusted for sex, race, chemistry, and BMI. Cell type comparisons were done in the form of (1) cell type *z* vs all other cell types to capture the global differences at *logFC* ≥ 0.585 and *FDR* ≥ 5% and (2) all pairwise comparisons among the endocrine cell types to detect genes that were uniquely upregulated in *z* at |*logFC*| ≥ 1 and *FDR* ≥ 5%. For glycemic states, we estimated all pairwise comparisons and reported differentially expressed genes at |*logFC*| ≥ 0.585 and *FDR* ≥ 5%. The functional characterization of the significant genes was done in *clusterProfiler v4.0*^115^ using the *enrichGO* function with background all genes of the GRCh38 genome. The enriched biological processes with set sizes in [10,500] were reported at FDR=5%.

### Differential Expression Analysis with a continuous covariate

We developed a Negative Binomial generalized linear model to identify genes whose expression levels varied as a function of a continuous covariate *X* such as Age and BMI. The significant genes were detected with the Likelihood Ratio (LR) test. For each gene *g*, the null model had the form: *E_g_* = *Y* + *Chemistry* + *Sex* + *Ancestry* where *E_g_* denotes gene’s *g* raw counts (pseudobulk) across islets of interest. The alternative model was *E_g_* = *sm.ns*(*X*,*df* = 2) + *Y* + *Chemistry* + *Sex* + *Race* of where *sm.ns* generated a basis matrix for natural cubic splines with 2 degrees of freedom, quantifying the smooth differences in mean expression as a function of the factor of interest *X*. Variable *Y* is any other factor to be adjusted and it was added here for convenience. For example, when looking for the genes whose expression differed across age (*X* factor), *Y* represented the BMI and reverse. The LR test assessed the goodness of fit of the two competing models based on the ratio of their likelihoods 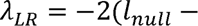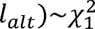 where *l_null_* and *l_alt_* are the log-likelihoods of the null and alternative models, respectively and λ*_LR_* is distributed as a chi-square with one degree of freedom. Rejection of the null model at significance level α was associated with a potentially significant finding. In practice, to minimize the chance of detecting genes with low p-values driven by a few outliers (over-smoothing due to the small sample size), we also estimate the Pearson correlation *rho_g_* between gene’s *g* CPM-normalized and log2-transformed data and *X*. The significant genes were detected at FDR=5% (λ*_LR_* test) and the empirical cutoff *rho_g_* ≥ 0.4.

### Cell Type Subclustering

The single-cell RNA-seq gene expression profiles of Alpha, Beta and Delta cell types were extracted and individually re-integrated as previously (see paragraph “*Data Integration*”). The UMAP was estimated from the first 20 PCs and the clustering was performed with the Leuven method and resolution parameter equal to 0.8. For each cell type, we compared gene’s *g* average expression in cluster *x_Z_* vs all other clusters by Seurat’s Negative Binomial (*negbinom*) model, adjusting for chemistry, sex, age and ancestry. We labeled genes as significant if.*logFC* ≥ 0.25 and *FDR* ≥ 1%.

### Cell culture

EndoC-βH3 cells were cultured in Advanced DMEM F-12 media (Invitrogen) containing BSA (SIGMA), Glutamax (Gibco), 2-beta mercaptoethanol (SIGMA), nicotinamide (SIGMA), sodium selenite (SIGMA), Penicillin/Streptomycin (Gibco) and Puromycin (Calbiochem) on ECM (SIGMA) and Fibronectin (SIGMA) coated flasks.

### Lentivirus production & transduction of cells

Plasmid pLKO-puro shRNA clones (Mission shRNA) were purchased from SIGMA. Lentivirus was produced in HEK293T cells co-expressing the shRNA plasmid together with psPAX2 packaging plasmid and pVSVG envelope plasmid. Virus was concentrated using Lenti-X Concentrator (Takara) and virus titer was quantified using p24 ELISA antigen assay (Takara). A MOI titer of 5 was used to transduce EndoC-βH3 cells at 1 ×10^6^ cells in culture media without puromycin. Media change to puromycin complete media was done 18hrs post transduction.

### RNA isolation

Total RNA was isolated from 3.5 ×10^5^ cells/sample 96hrs post transduction. Cells were collected for RNA extraction using TRIZOL (Invitrogen), phase separation was achieved using chloroform. Isopropanol was used for RNA precipitation using glycogen as a carrier, the pellets were washed using 75% ethanol, air-dried, and resuspended in DEPC water. RNA was measured using Nanodrop. Total RNA was used to perform qPCR using RNA to CT kit (Invitrogen) and FAM-Taqman probes (Invitrogen) and analyzed on QuantStudio 7 (Applied Biosystems) normalized to *ACTNB* Taqman probe.

### Insulin secretion assay

EndoC-βH3 cells infected with the lentivirus were seeded onto coated 24 well plates at 1.75 ×10^5^ cells/well. 72 hrs post transduction the cells were incubated overnight in Starvation media [DMEM no glucose, containing BSA (SIGMA), human Transferrin (SIGMA), Glutamax (Gibco), 2-beta mercaptoethanol (SIGMA), nicotinamide (SIGMA), sodium selenite (SIGMA)]. After 18h cells were equilibrated in KRBH buffer containing no glucose for 1 hour, before stimulated insulin secretion was measured by static incubation of KRBH buffer containing 0mM and 20mM Glucose for 1 hour. The supernatant was collected and stored at −20°C until human insulin ELISA (Mercodia). After glucose stimulation, KRBH buffer was collected and the cells were lysed with TETG solution [1M Tris pH 8.0, Triton X-100, Glycerol, 5M NaCl, 0.2M EGTA, and distilled water along with 1X cOmplete, protease inhibitor cocktail (Roche)]. The lysate was centrifuged at 3,000 rpm for 5 minutes and stored at −20°C until human insulin ELISA (Mercodia) according to manufacturer’s instruction. Total protein was measured using BCA kit (Thermo Fisher) and insulin secretion and content normalized to total protein content per sample.

### Flow cytometry

EndoC-βH3 cells infected with the lentivirus were seeded onto coated 24 well plates at 1.75 ×10^5^ cells/well. 90 hrs post transduction cells were collected from the plate using Trypsin (Gibco) and stained using FITC Annexin V Apoptosis Detection Kit with 7-AAD (BioLegend) according to Manufacturer’s instruction. The samples were run on Fortessa (BD Sciences), and data was analyzed via FlowJo Software (BD Sciences).

### Functional annotation of DEGs

We obtained the summary statistics of all genetic variants significantly associated with T2D at genome-wide significance *P*L<L5L×10^−8^ from multiple ancestry metanalyses - T2DGGI^37^, DIAMANTE^38^, MVP^39^ and AGEN^40^. For each T2D associated variant, we identified proxy variants linked at r^2^≥0.80 across all ancestries based on 1000Genomes Phase 3 data. To determine if these T2D variants serve as eQTLs for our identified DEGs, we downloaded summary statistics data of *cis*-eQTL associations from TIGER^41^ atlas (https://tiger.bsc.es/), which includes 404 human pancreatic islet samples of European descent and reports >1.11 million significant eQTLs in >21,115 eGenes at 5% FDR. We retrieved all variants reported as eQTLs for our identified DEGs at p-value < 0.05, with a consistent direction of association across all four independent cohorts (++++ or ----) in the TIGER dataset. We then cross-referenced the T2D genetic variant list and the eQTL list and identified eQTL variants for 41/511 DEGs that were also associated with T2D genetic risk.

Protein-level association data in T2D vs. ND whole islets was downloaded from the HumanIslets^56^ consortium (https://www.humanislets.com/#/) that reports mass spectrometry (MS)-based bulk protein expression data in 300 hand-picked islets per sample. P values were calculated by multiple linear regression and adjusted for multiple testing with FDR.

Phenotypic data from whole gene knockout mouse lines was obtained from the International Mouse Phenotyping Consortium^62^ web portal (https://www.mousephenotype.org/). The latest data release (Nov 2024) reports 9,073 phenotyped genes, with knockout mice produced and characterized by various institutional members of IMPC. All mice used in IMPC studies have a C57BL/6N genetic background, with supporting mice derived from C57BL/6NJ, C57BL/6NTac or C57BL/6NCrl strains. Glucose tolerance was assessed by initial response to glucose challenge and/or calculating the area under the glucose response curve from an intraperitoneal glucose tolerance test (IPGTT).

---

**Supplementary Figure 1:**
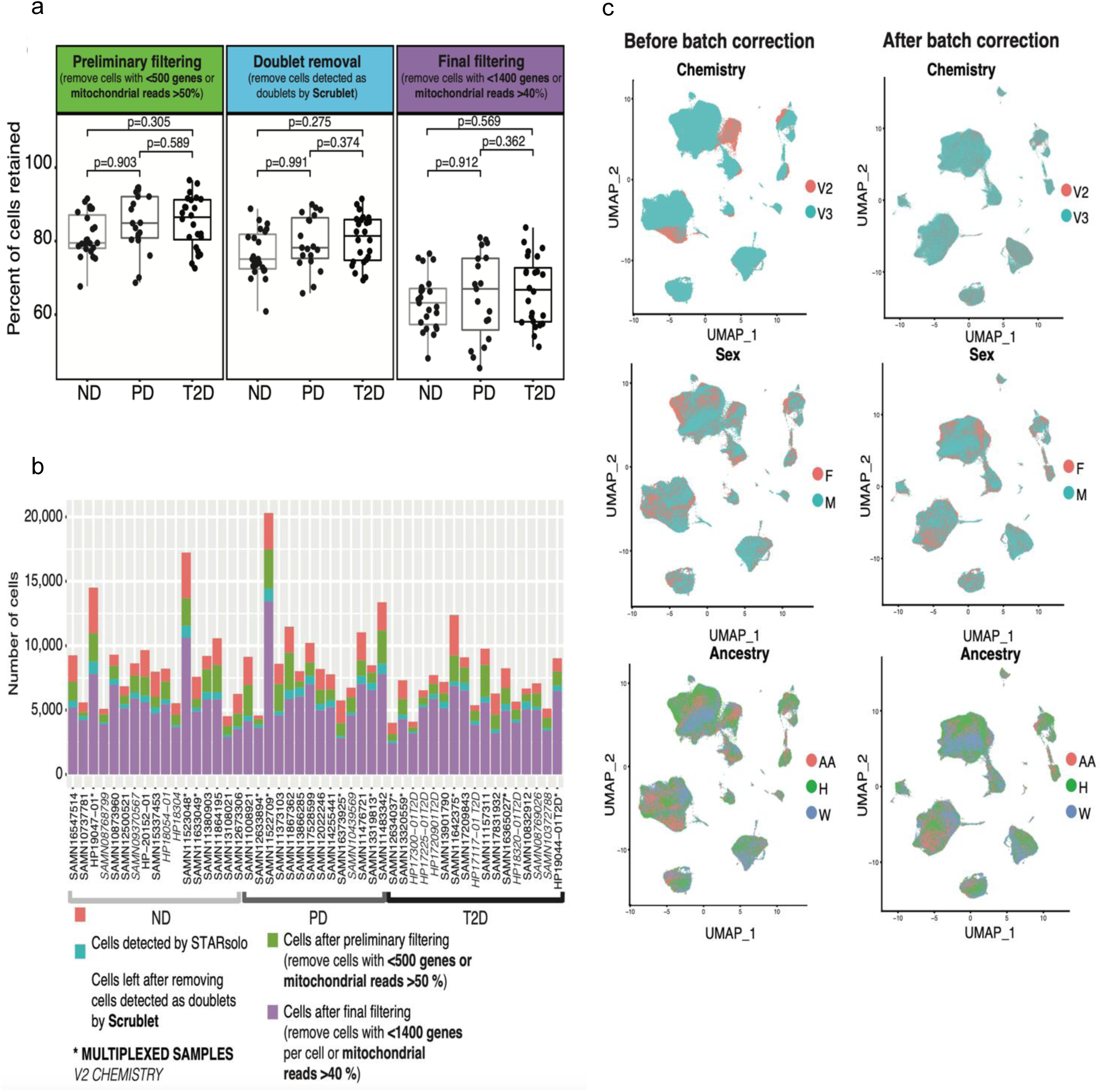
Quality control measures of single cell transcriptomes from each donor. **(a)** Percent of cells retained (y-axis) for non-diabetic (ND), prediabetic (PD) or type 2 diabetic (T2D) donors after each filtering step (x-axis); each dot represents a donor. Bonferroni-adjusted p-values from Tukey’s honest significance test are shown. **(b)** Stacked bar plot indicating number of cells retained (y-axis) per donor (x-axis) after each QC filtering step. Samples with multiplexed runs are typed in bold; 10X sequencing v2 chemistry runs are italicized. **(c)** UMAPs of single cell transcriptomes after preliminary filtering before and after batch correction for sequencing chemistry, sex, and ancestry.

**Supplementary Figure 2:**
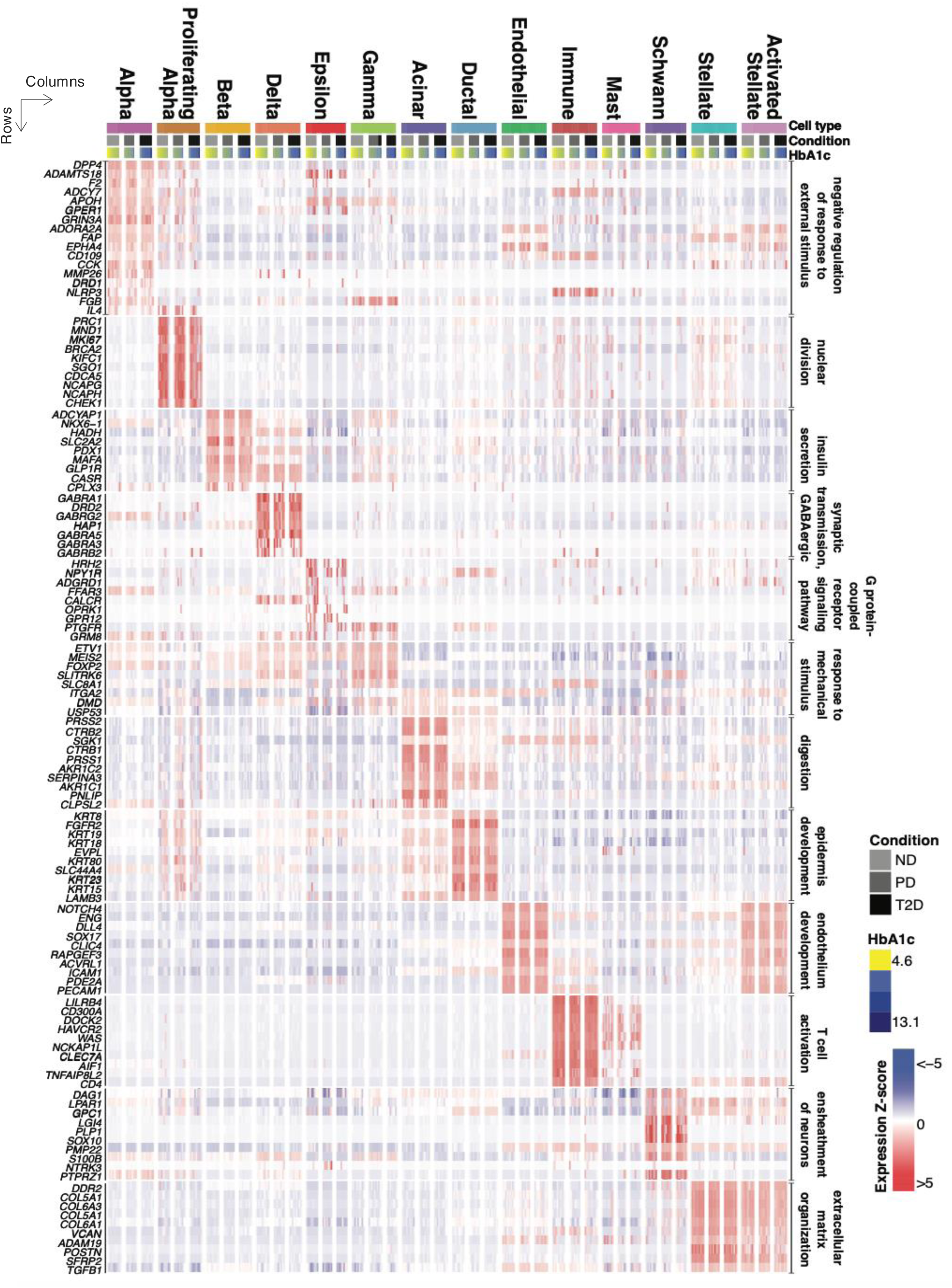
Heatmap of aggregated (pseudobulk) marker gene expression (left; rows) representing enriched GO terms (right) for each islet cell type (columns). Individual islet donors are grouped by glycemic status (grayscale) and sorted from lowest to highest reported HbA1c levels (yellow-blue gradient) for each cell type and glycemic status shown. ND: Non-diabetic, PD: prediabetic, T2D: Type 2 diabetic. Full set of enriched marker genes and GO terms for each cell type are provided in Supplementary Table 3 and 4.

**Supplementary Figure 3:**
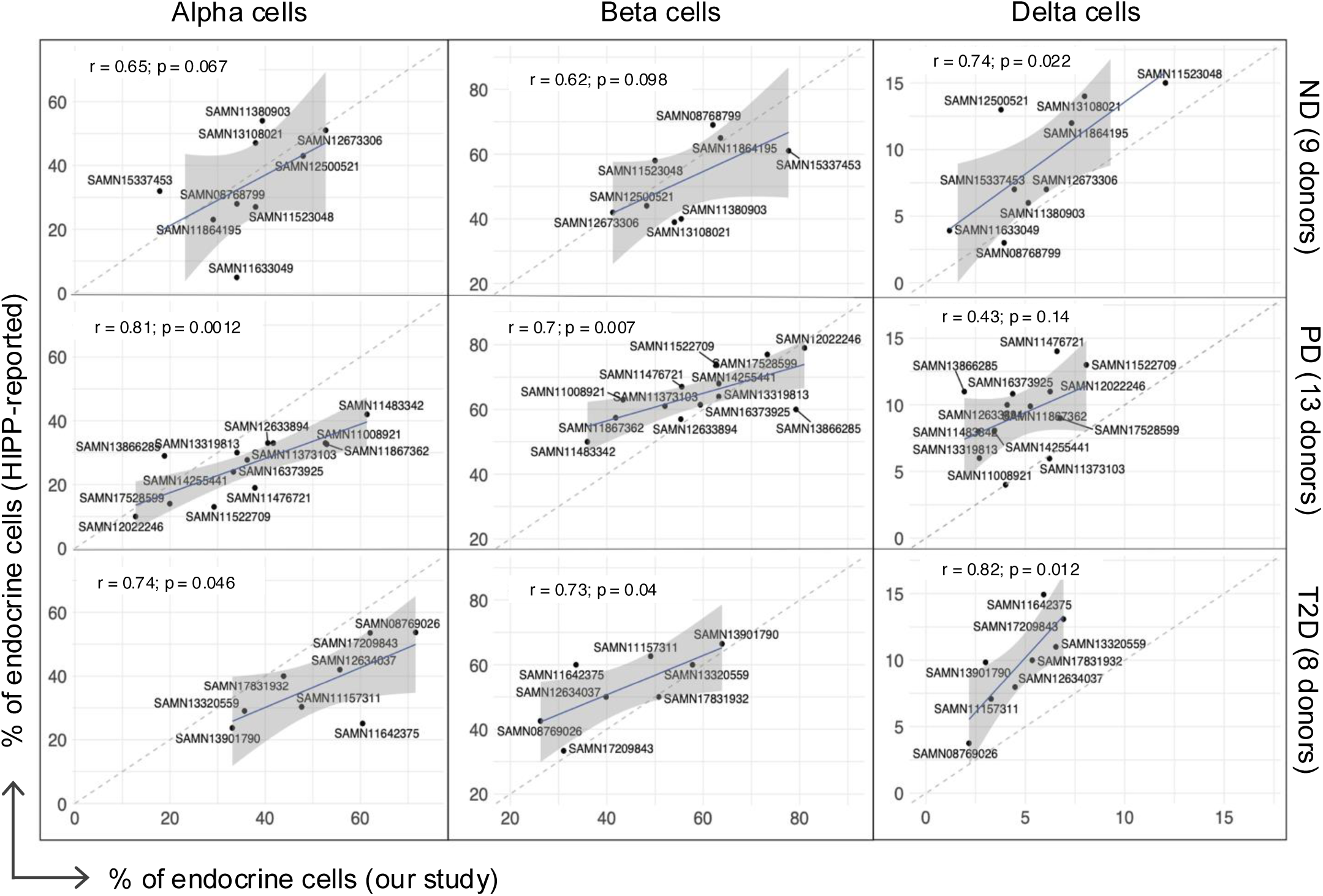
Scatter plots of spearman correlation (r) between alpha, beta, and delta cell endocrine proportions reported by the the Human Islet Phenotyping Program (HIPP, y-axis) for 9 Non-diabetic (ND), 13 prediabetic (PD), and 8 type 2 diabetic donors and as detected by our scRNA-seq counts (x-axis) from the corresponding donors. The identity line (dashed) and the line of fitted linear regression model (solid blue) are shown.

**Supplementary Figure 4:**
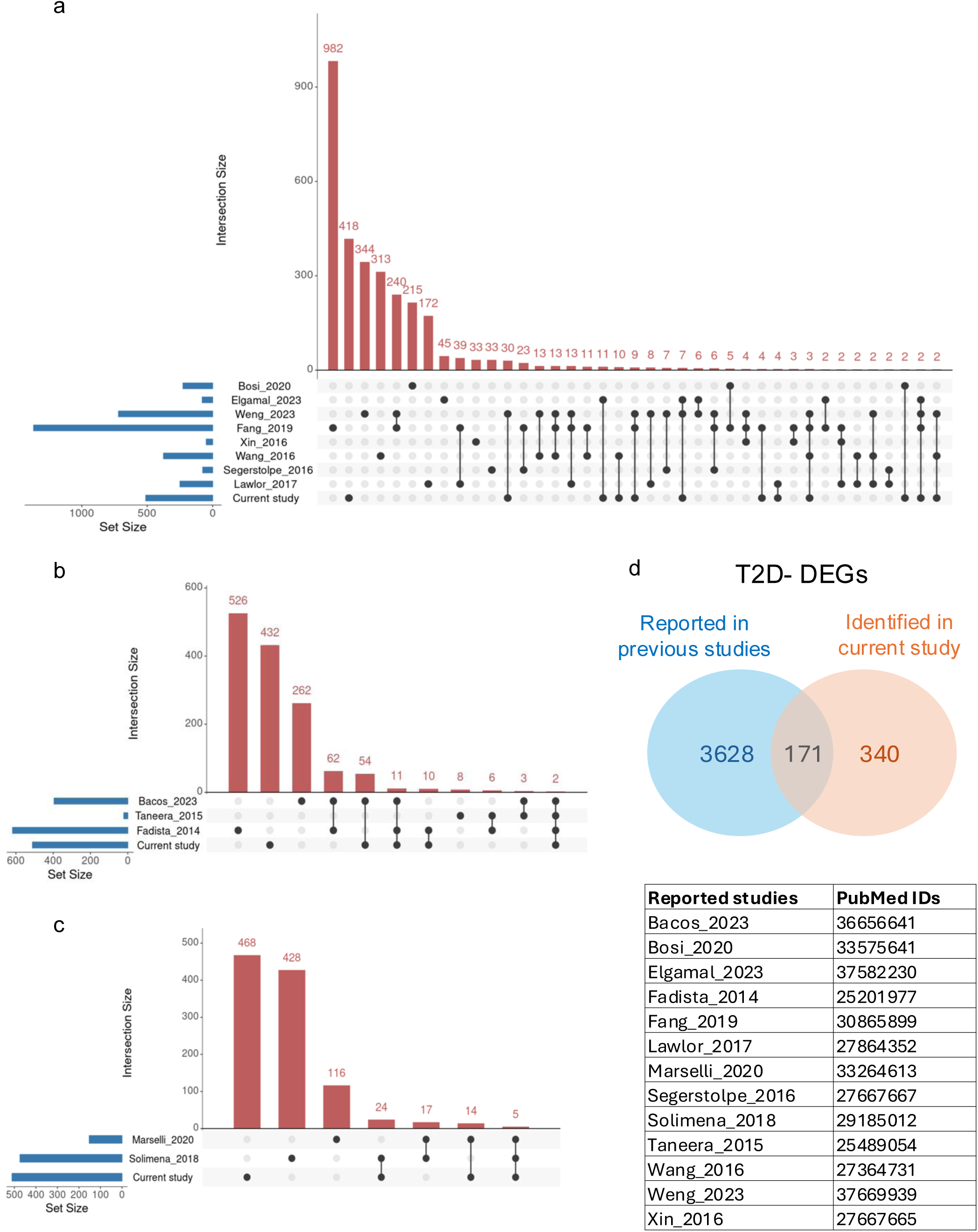
UpSet plots showing overlap between the identified T2D vs ND β-cell differentially expressed genes (DEGs) with previously reported DEGs – β-cell scRNA-seq **(a)**, islet RNA-seq **(b),** and sorted β-cell RNA-seq **(c)**. Overlaps containing at least 2 common genes are shown. **(d)** Venn diagram showing our replication of 171 previously reported genes and identification of 340 novel T2D-DEGs in current study.

**Supplementary Figure 5:**
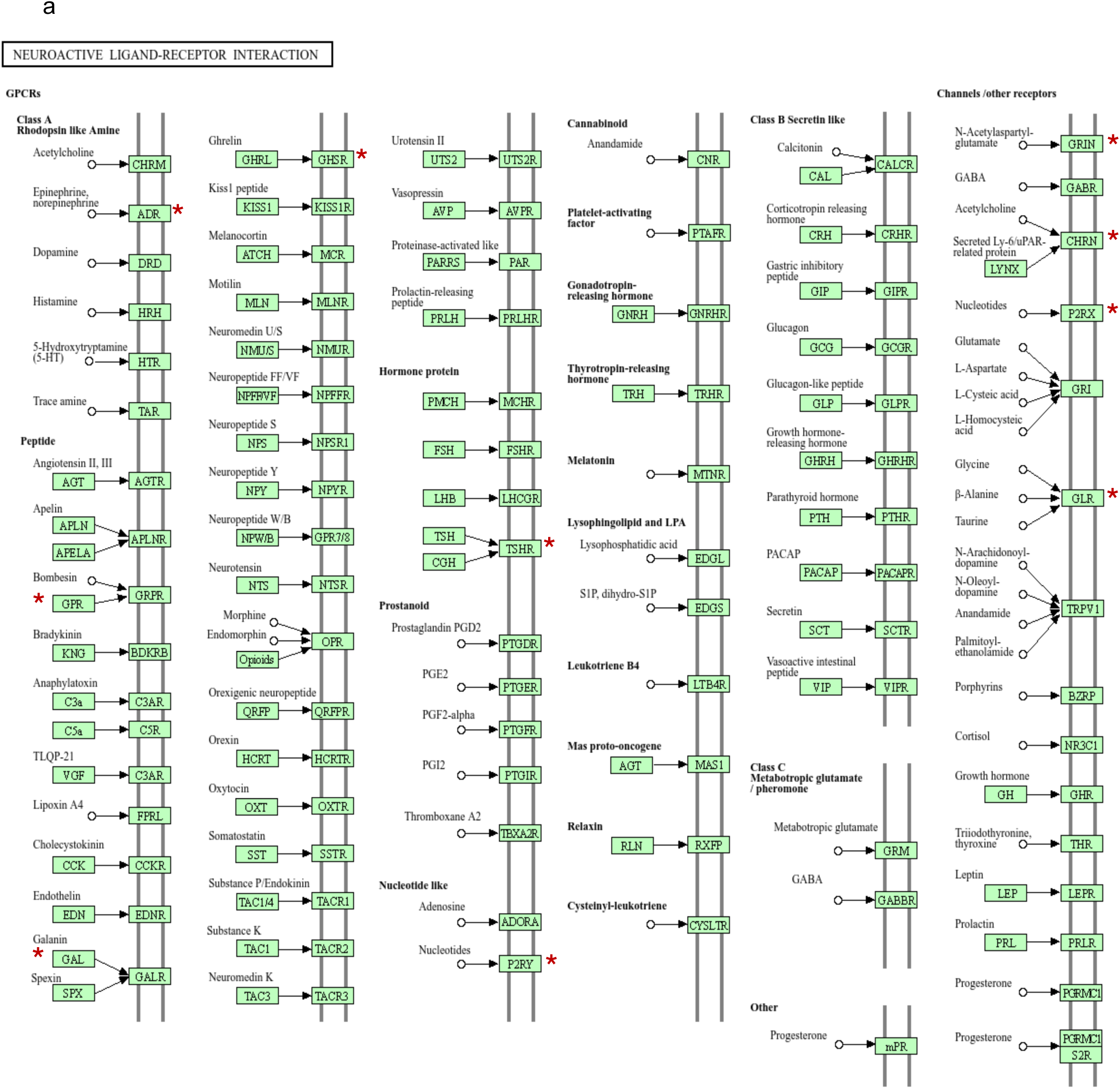
Primary pathways associated with T2D β-cell differentially expressed genes (DEGs). **(a)** ‘Neuroactive ligand receptor interaction’ enrichment in upregulated genes. **\*** marks identified T2D β-cell DEGs in the pathways. Image sources: KEGG and Wikipathways.

**Supplementary Figure 5(b).**
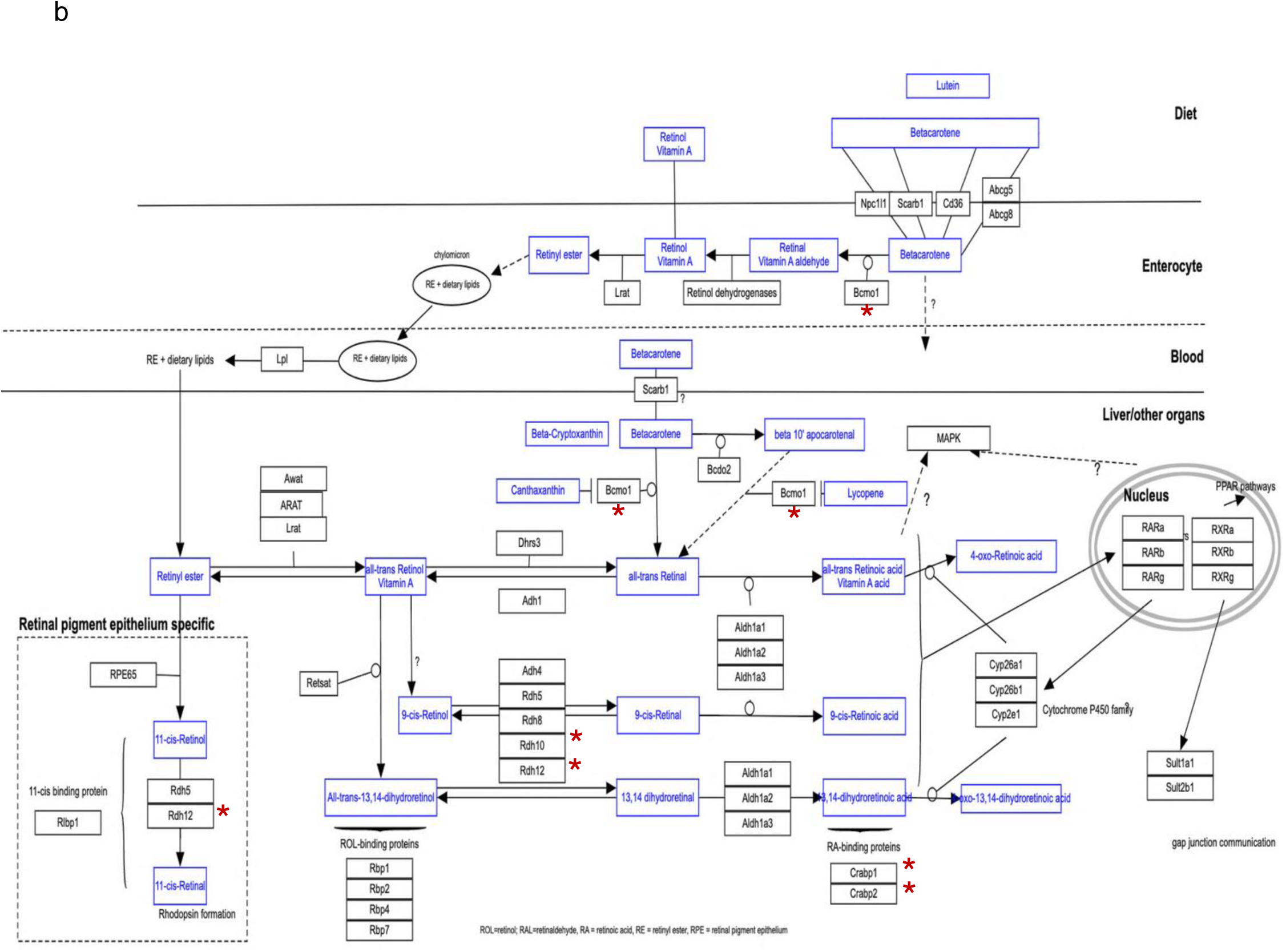
‘Vitamin A and carotenoid metabolism’ in downregulated genes. **\*** marks identified T2D β-cell DEGs in the pathways. Image sources: KEGG and Wikipathways.

**Supplementary Figure 6:**
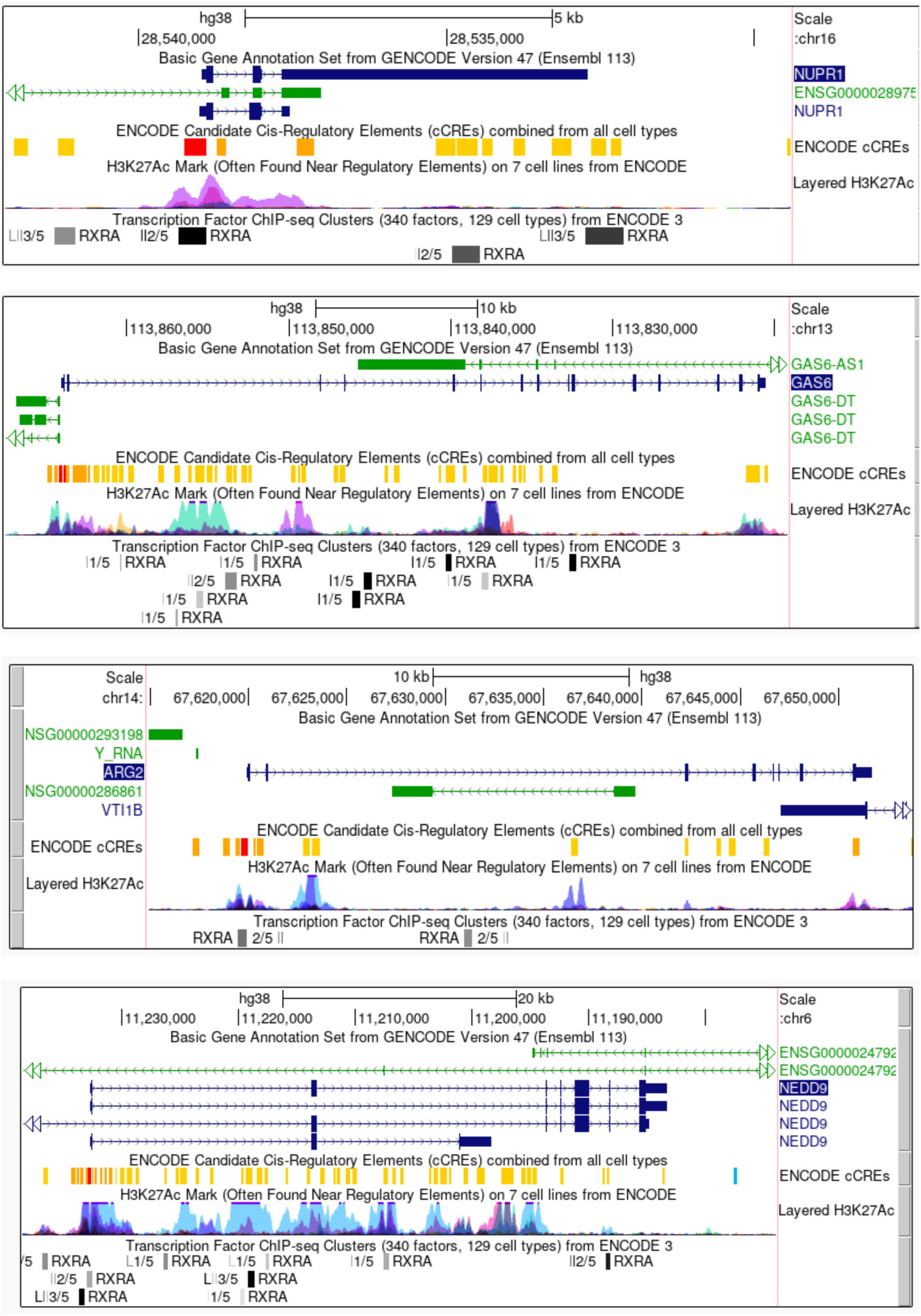
Genomic locus snapshots of β-cell death-related DEGs, highlighting the presence of retinoic acid (RA) response elements and RA receptor (RXRA) binding. These elements regulate the expression of RA target genes. Data source: ENCODE TF ChIp-seq, UCSC Genome Browser.

**Supplementary Figure 7:**
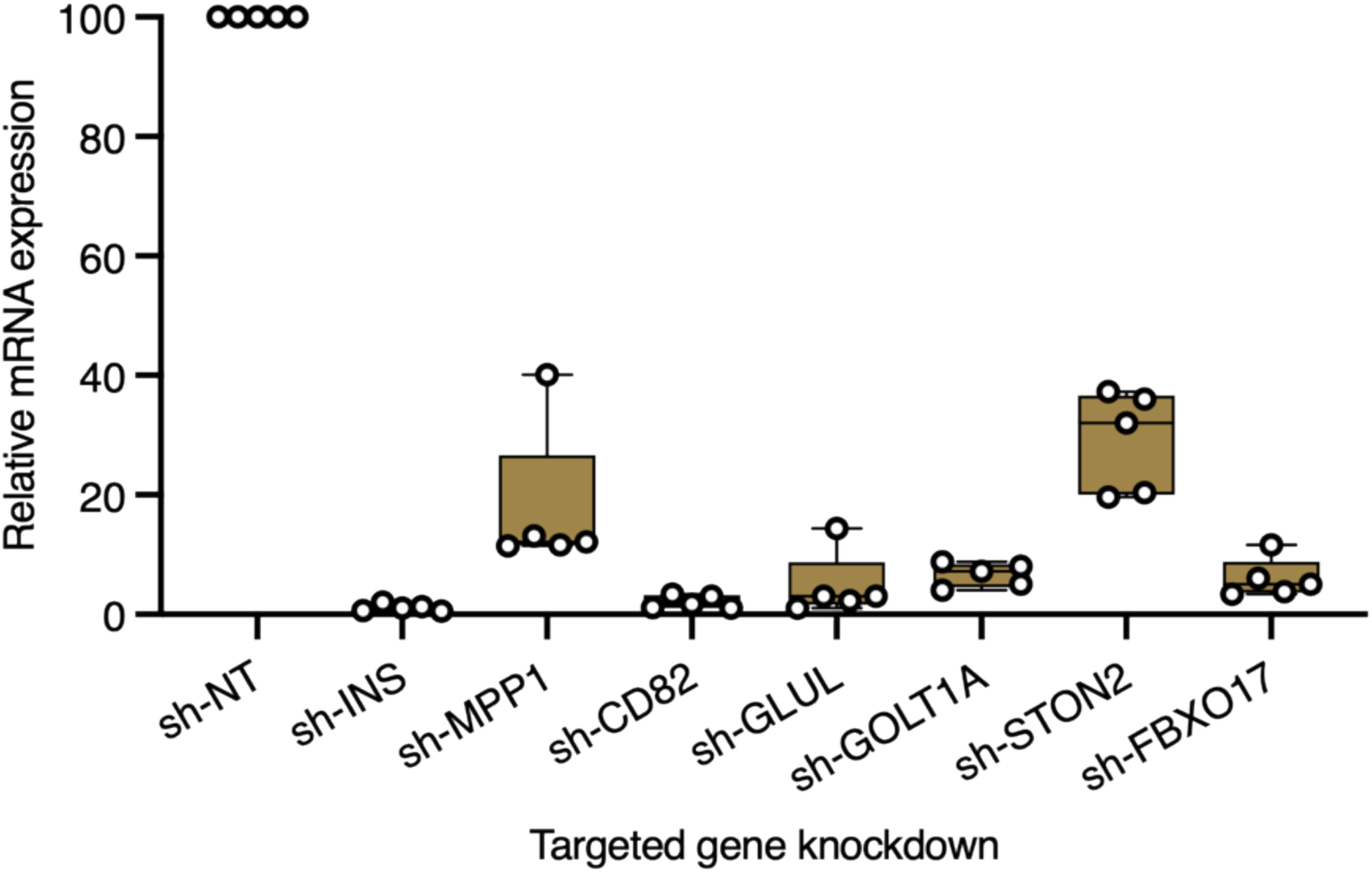
shRNA mediated knockdown of selected T2D-downregulated β-cell genes in human EndoC-βH3 cells. Gene expression is shown relative to that in non-targeting (NT) shRNA transduced cells. *INS* was tested as a positive control. Whiskers represent minimum and maximum knockdown efficiencies from 5 independent experiments.

**Supplementary Figure 8:**
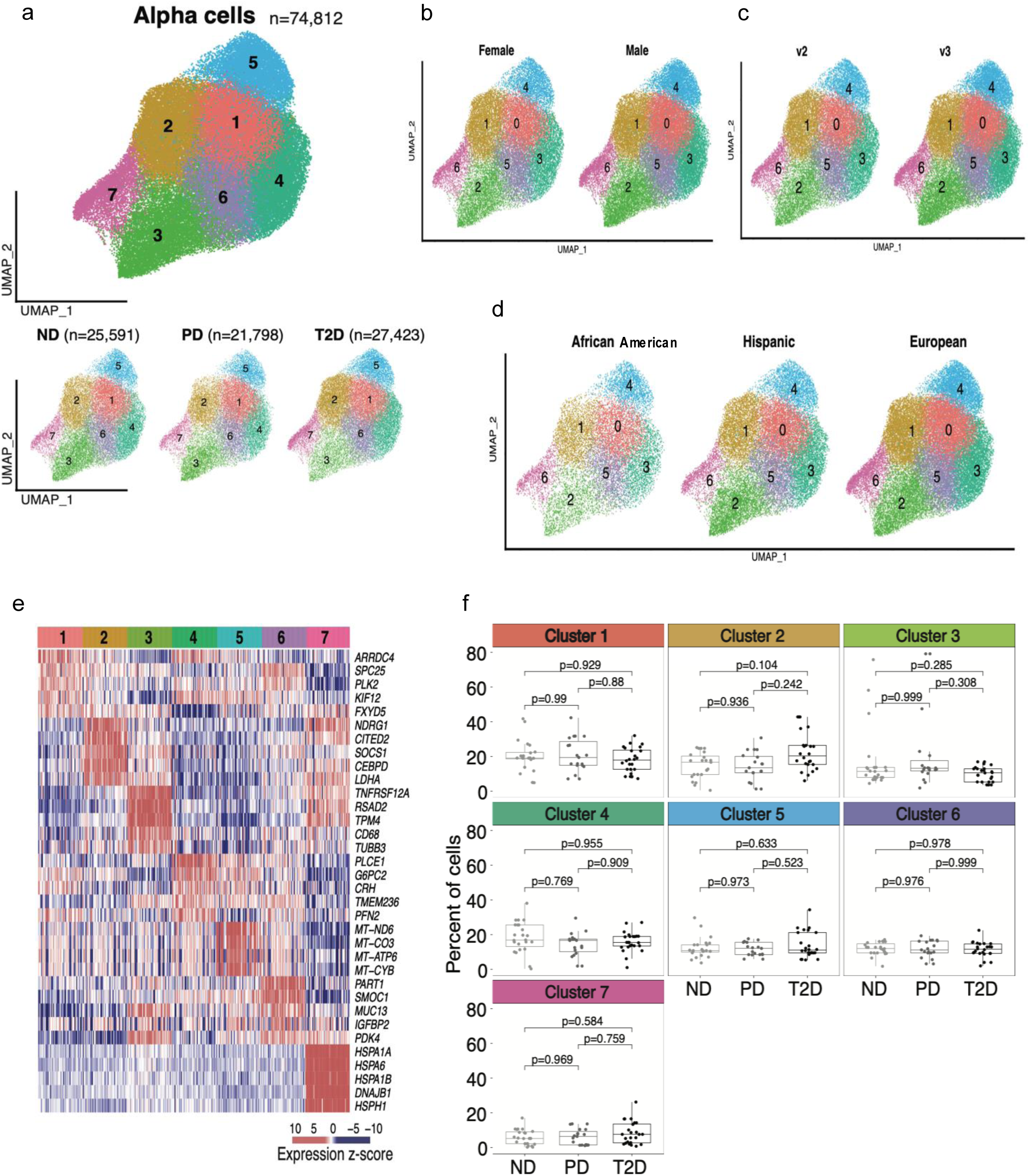
UMAPs of alpha-cell subpopulations shown for each glycemic status (**a)**, sex (**b**), scRNA sequencing chemistry (**c**; v2 or v3), and self-reported ancestry (**d**; European, African American and Hispanic). Number of cells per cluster (n) is indicated in parentheses. (**e**) Heatmap of normalized marker gene expression in alpha-cell subpopulations. (**f**) Putative ⍺-cell subpopulation proportions in non-diabetic (ND; n = 17), prediabetic (PD; n = 14) and type 2 diabetic (T2D; n = 17) donors. Individual dots display per-donor proportions in each group. P-values between groups are calculated from Tukey’s honestly significance test, adjusting for Bonferroni correction.

**Supplementary Figure 9:**
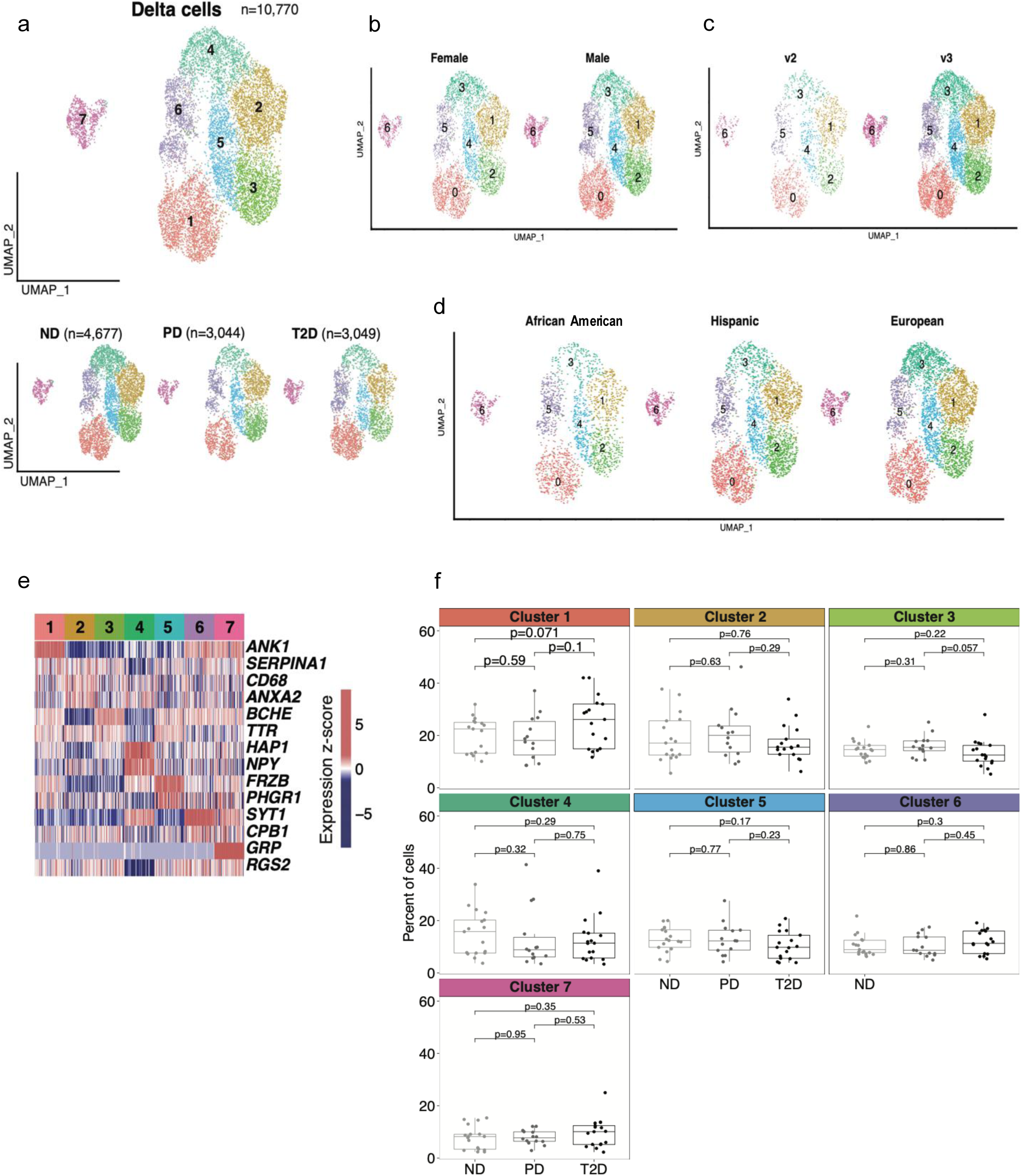
UMAPs of delta-cell subpopulations shown for each glycemic status (**a)**, sex (**b**), scRNA sequencing chemistry (**c**; v2 or v3), and self-reported ancestry (**d**; European, African American and Hispanic). Number of cells per cluster (n) is indicated in parentheses. (**e**) Heatmap of normalized marker gene expression in delta-cell subpopulations. (**f**) Putative α-cell subpopulation proportions in non-diabetic (ND; n = 17), prediabetic (PD; n = 14) and type 2 diabetic (T2D; n = 17) donors. Individual dots display per-donor proportions in each group. P-values between groups are calculated from Tukey’s honestly significance test, adjusting for Bonferroni correction.

**Supplementary Figure 10:**
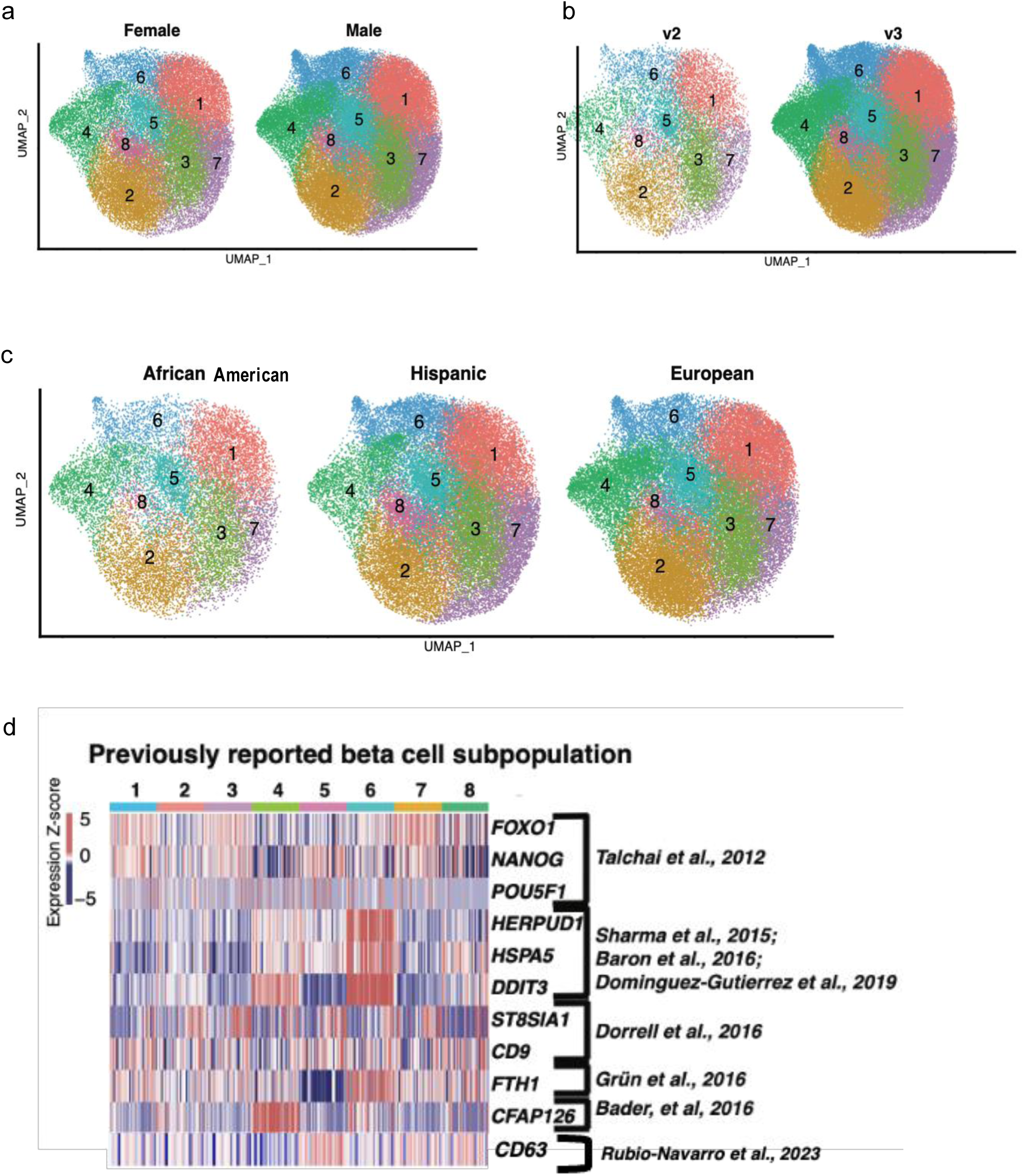
**a)** UMAPs of putative beta-cell subpopulations shown for each sex (**a**), scRNA sequencing chemistry (**b**; v2 and v3), and self-reported ancestry (**c**; European, African American and Hispanic). **d)** Heatmap of scaled expression of previously reported beta-cell subpopulation marker genes^72,74–78,95,96^ in this study.

## Notes

https://cellxgene.cziscience.com/collections/58e85c2f-d52e-4c19-8393-b854b84d516e

https://thejacksonlaboratory.shinyapps.io/TAPIC_Stitzel_Lab/

https://zenodo.org/records/14656366

### Competing Interest Statement

The authors have declared no competing interest.

